# *N*-linked glycans confer protein stability and modulate multidrug efflux pump assembly *in Campylobacter jejuni*

**DOI:** 10.1101/585463

**Authors:** Sherif Abouelhadid, John Raynes, Tam T.T. Bui, Jon Cuccui, Brendan W. Wren

## Abstract

It is now apparent that nearly all bacteria species have at least a single glycosylation system, but the direct effect(s) of these protein post translational modifications are unresolved. In this study, we used the general *N*-linked glycosylation pathway from *Campylobacter jejuni* to investigate the biophysical roles of protein modification on the CmeABC multidrug efflux pump complex. The study reveals the multifunctional role of *N*-linked glycans in enhancing protein thermostability, stabilising protein complexes and the promotion of protein-protein interaction. Our findings demonstrate, for the first time, that regardless of glycan diversification among domains of life, *N*-linked glycans confer a common evolutionary intrinsic role.

## Introduction

Glycosylation, the process of covalently attaching a glycan (mono-, oligo-, or polysaccharide) to an amino acid side chain, is the most prevalent post-translational modification found on proteins, occurring in all domains of life. The attachment of a carbohydrate moiety to certain amino acid side chains in proteins was traditionally thought to be exclusive to eukaryotes and archaea. In the last decade, protein glycosylation, both *N*- and *O*-linked, has been increasingly reported in pathogenic bacteria such as well as commensal bacteria^1^. Functional analysis of the role of *N*-glycans in eukaryotes shows multiple roles. Intrinsically, glycans confer protein stabilisation through a plethora of mechanisms such as, promoting secondary structure formation, accelerating protein folding, slowing the unfolding rate^2^, enhancing protein thermostability^3^ and reducing aggregation^4^. Extrinsically, *N*-linked glycosylation modulates protein-protein interactions and protein targeting^5^. Notably, prokaryotes and eukaryotes share similar biosynthetic pathways of general *N*-linked glycosylation, indicating a common evolutionary pathway^6^. However, the role of *N*-linked glycans remains poorly studied in prokaryotes.

The first characterisation of an *N*-linked glycosylation in a bacterium was in the enteric pathogen *Campylobacter jejuni* which now has one of the most studied of all prokaryotic glycosylation pathways^7^. Genomic and proteomic studies have demonstrated that, PglB is an oligosaccharyltransferase responsible for the covalent attachment of *N*-linked glycan (GalNAc -α1,4-GalNAc-α1,4-GalNAc-[Glcβ1,3-]GalNAc-α1,4-GalNAc-α1,4-GalNAc- α1,3-Bac-β1; where GalNAc is *N*- acetylgalactosamine; Glc is glucose; diBacNAc is 2,4- diacetamido −2,4,6- trideoxyglucopyronose) to the asparagine residue in the acceptor sequon D/E-X_1_-N-X_2_- S/T where X_1_ and X_2_ are any amino acid except proline^7^. In depth studies on *N*-linked glycan diversity have revealed that there are 16 different glycans present in *Campylobacter* genus. Surprisingly, the first two glycans (diBacNAc and HexNAc) were conserved in all species^8, 9^. Whilst the role of *N*-linked glycan remains to be elucidated, disruption of the *N*-linked glycosylation pathway resulted in pleotropic effects such as decreased chicken colonisation^10^, reduced adherence to intestinal cells^11^ as well as impaired bacterial competence^12^. These studies did not ascertain the direct role of *N*- linked glycans in the *C. jejuni* glycoproteome. To address this, we sought to investigate the biophysical role of *N*-linked glycosylation on a representative post translationally modified glycoprotein. We demonstrate a critical role for glycosylation by focusing on a resistance-nodulation-division type (RND) multidrug efflux pump denoted *<u>C</u>ampylobacter* <u>m</u>ultidrug <u>e</u>fflux; CmeABC.

CmeABC is a tripartite molecular assembly of glycoproteins, CmeB; an inner membrane multidrug transport protein, CmeA; a periplasmic fusion protein and CmeC; an outer membrane associated channel. The complex plays a key role in chicken colonization^13^, as well as being responsible for multidrug resistance (MDR)^14^. Previously, we demonstrated that disrupting *pglB* in *C. jejuni* impaired the efflux activity of CmeABC resulting in significantly higher ethidium bromide accumulation when compared to the wildtype (Abouelhadid S, *et al* submitted manuscript). The absence of glycosylation on the CmeABC locus within *C. jejuni* was also shown to reduce resistance to four different antibiotic classes. Here in, we show that the loss of *N*-linked glycans in CmeABC is the sole reason for this phenotype and not a pleiotropic effect caused by *pglB* disruption. We also unravel the intrinsic role of *N*-linked glycans in a) modulating global protein structure b) enhancing glycosylated CmeA; g2CmeA thermostability and c) significantly slowing the unfolding rate of g2CmeA. Finally, we evaluate the extrinsic role of *N*-linked glycans in the molecular assembly of CmeABC to discern the difference in binding kinetics of CmeA variants to CmeC. The study highlights the multifunctional role of *N*-linked glycans in enhancing protein thermostability, stabilising protein complexes and the promotion of protein-protein interaction. Here we present a model N-linked glycosylation system with a tractable phenotype to be used in studying glycan evolution, function and diversity. Our findings demonstrate, for the first time, that regardless of glycan diversification among domains of life, *N*-linked glycans seem to confer a common evolutionary intrinsic role.

## Results

### *N*-linked glycans do not affect CmeABC protein expression nor protect CmeC from native proteolytic degradation

Scrutiny of the *C. jejuni* NCTC11168 genome revealed the presence of 13 multidrug efflux transporters, which appear conserved in the species^15^. Genetic and biochemical testing demonstrated that *cmeABC* is located in an operon and encodes the predominant multidrug efflux pump in *C. jejuni*. In addition to its central role in extruding structurally non-related compounds such as antimicrobials, bile salts, dyes and heavy metals^14^, CmeABC has been reported to function interactively with CmeDEF^16^; a secondary multidrug efflux pump in *C. jejuni*. Glycoproteomic analysis of *C. jejuni* demonstrated that the CmeABC complex is multi *N*-linked glycosylated where CmeA is glycosylated at positions ^121^DF<u>N</u>R^124^ and ^271^DNNNS^275^, CmeB is glycosylated at position ^634^DR<u>N</u>VS^648^ and CmeC is glycosylated at position ^47^ET<u>N</u>SS^51^. Notably, CmeE has also been shown to be *N*-glycosylated^17^. Previously, we showed that the multidrug efflux pump is impaired in the glycosylation deficient *C. jejuni*. This resulted in a significant increase in ethidium bromide accumulation more than the parent strain and a reduction in antibiotics resistance (Abouelhadid S, *et al* submitted manuscript). We hypothesise that this deficiency may be due to assembly destabilisation as a consequence of glycosylation aberration. To address this question, we sought to study the major multidrug efflux pump of *C. jejuni*, in a glycosylation deficient CmeABC complex. We conducted the experiments in a *C. jejuni cmeD*::*cat*^r^ background strain in order to avoid the interaction of CmeDEF with CmeABC that might mask the functional role of *N*-linked glycans. This parent strain was used to construct a *C. jejuni* WTCmeABC strain and a *C. jejuni* glycosylation altered strain; g0CmeABC whereby, N->Q in each glycosylation sequon (D/E-X_1_-N-X_2_- S/T where X_1_ and X_2_ are any amino acids other than proline). Since *cmeABC* is an operon, we added a 6x His tag at the C-terminal of CmeC to monitor changes in CmeABC expression. We then grew both WTCmeABC and g0CmeABC strains to an OD_600_ =0.4-0.5, tetracycline was then added to inhibit protein synthesis. CmeC levels in both strains were monitored for 2 hours. Our data show no difference in CmeC expression in WTCmeABC and g0CmeABC strains Fig 1, A. Once inserted in the outer membrane CmeC might be unreachable region to proteases. Therefore, we conclude that *N*-linked glycans, in this case, do not protect CmeC from native proteolytic degradation. We observed two bands in the WTCmeC (lanes 2, 4, 6, 8) that migrated slower than the band corresponding to non-glycosylated CmeC in g0CmeC (lanes 1, 3, 5, 7). Western blot analysis confirmed that the two bands observed in WTCmeC lanes correspond to two glycospecies for the protein. Fig 1, B. Our bioinformatic analysis of the primary amino acid sequence shows that CmeC has two glycosylation sequons; ^30^EANYS^34^ and ^47^ETNSS^51^ in *C. jejuni* NCTC11168. Structurally both of the glycosylation sequons are located in flexible loops^18^ Fig 6, B.

**Fig 1.**
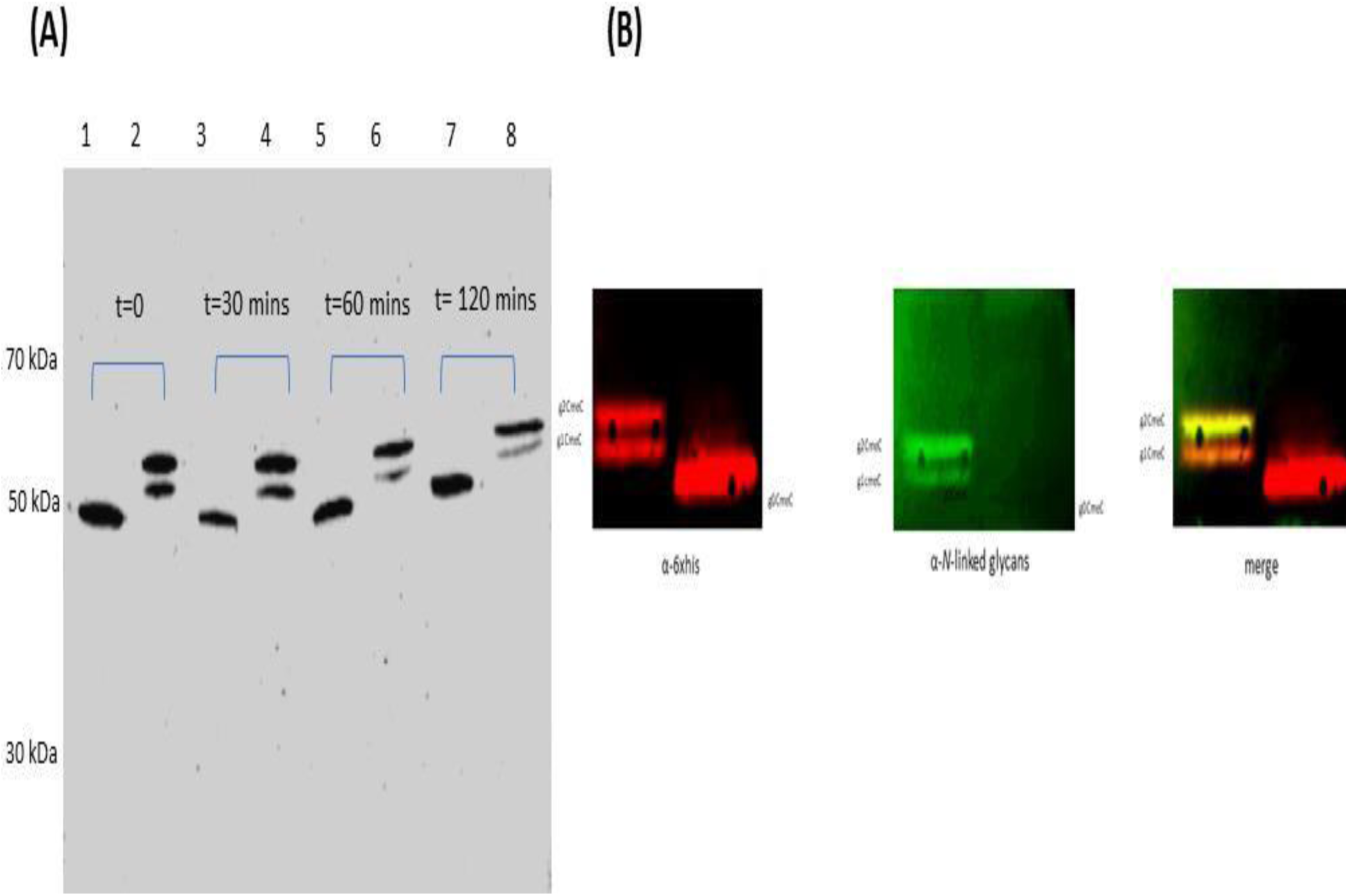
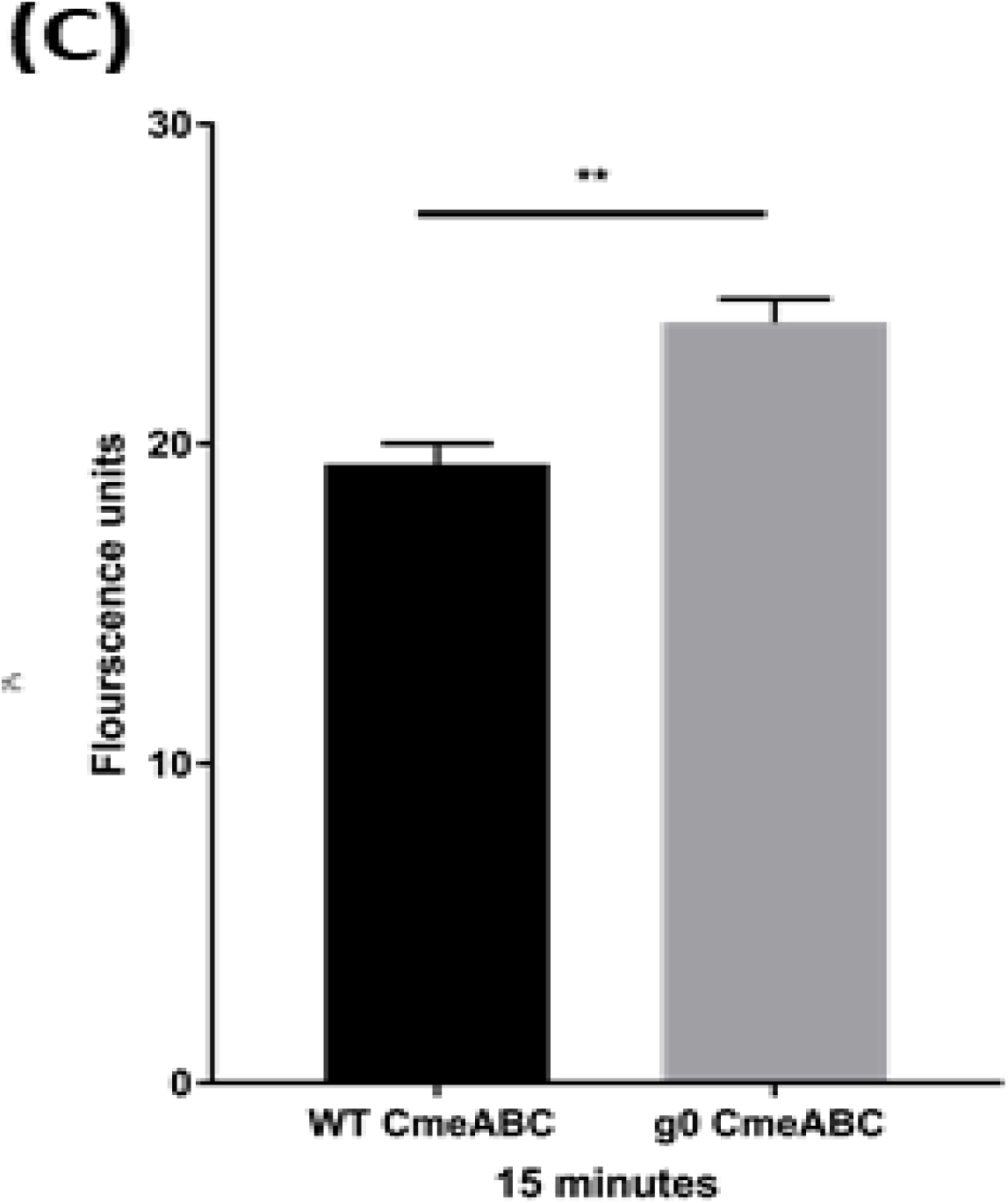
Functional studies and effect of glycosylation on WTCmeABC and g0CmeABC (A) Proteolytic degradation assay of CmeC and gCmeC. Cells were grown to OD_600_ =0.4 then Chloramphenicol and Tetracycline were added. Initial sample were withdrawn and labeled t=0, then samples were taken every 30 minutes. Cells were stored on ice, centrifuged, lysed by sonication then incubated with 2% SDS and Sodium sarkosyl for 2 hours at room temperature. Cells debris were then pelleted by centrifugation and supernatants were mixed 1:1 with Laemmli loading buffer supplemented with DTT. Proteins were then separated by SDS-PAGE followed by electroblotting to PVDF membrane, 6xhis tagged CmeC was probed by 1ry anti-6xhis mouse antibody and visualized by Li-COR odyssey. Equal amount of proteins was loaded, lane 1, 3, 5, 7, *C. jejuni* g0CmeC; lane 2, 4, 6, 8 *C. jejuni* WTCmeABC (B) Western blot detection of CmeC variants, lane 1, g2CmeC; lane 2, g0CmeC. Proteins were detected with anti-6xHis antibody and *N*-linked glycans were detected with SBA lectin-biotin. (C) Ethidium bromide accumulation test in *C. jejuni* strains. 30 ml Brucella broth was separately inoculated with overnight culture of *C. jejuni* WTcmeABC (black) and *C. jejuni* g0CmeABC (grey) to O.D_600_ 0.1. Cells were growntill O.D_600_ 0.4-0.5 then spun down, washed and resusupended to OD_600_ 0.2 in 10 mM sodium phosphate buffer pH 7. Cells were then incubated in VAIN for 15 minutes at 37°c then Ethidium bromide was added to final concentration of 0.2 mg/ml. Fluorescence was read at excitation and emission for 20 minutes at 37°c accumulation in *C. jejuni* strains at 15 minutes. The data represents the mean of three biological replicates, two technical replicates each. Significance was calculated using Mann-Whitney test. \*\**P*<0.005

### *N*-linked glycans affect multidrug efflux pump efficiency

To examine the role of *N*-linked glycans in CmeABC molecular assembly, we assessed the efficiency of the multidrug efflux pump using an ethidium bromide accumulation assay. Ethidium bromide accumulation was 22% higher in g0CmeABC when compared to WTCmeABC. This difference was consistent at 5, 10 and 15 minutes, indicating an impairment in the extrusion of ethidium bromide from g0CmeABC Fig 1, C. To confirm this finding, E-test antibiotic strips were used to calculate the minimum inhibitory concentration (MIC) of four non-structurally related antibiotics that have different mechanisms of actions. In comparison to WTCmeABC, an increase in antibiotic susceptibility was noticed in g0CmeABC confirming the previous finding Table 1. The results indicate that *N*-linked glycans play a role in enabling the multidrug efflux pump to work efficiently in the *C. jejuni* cell.

**Table 1.**
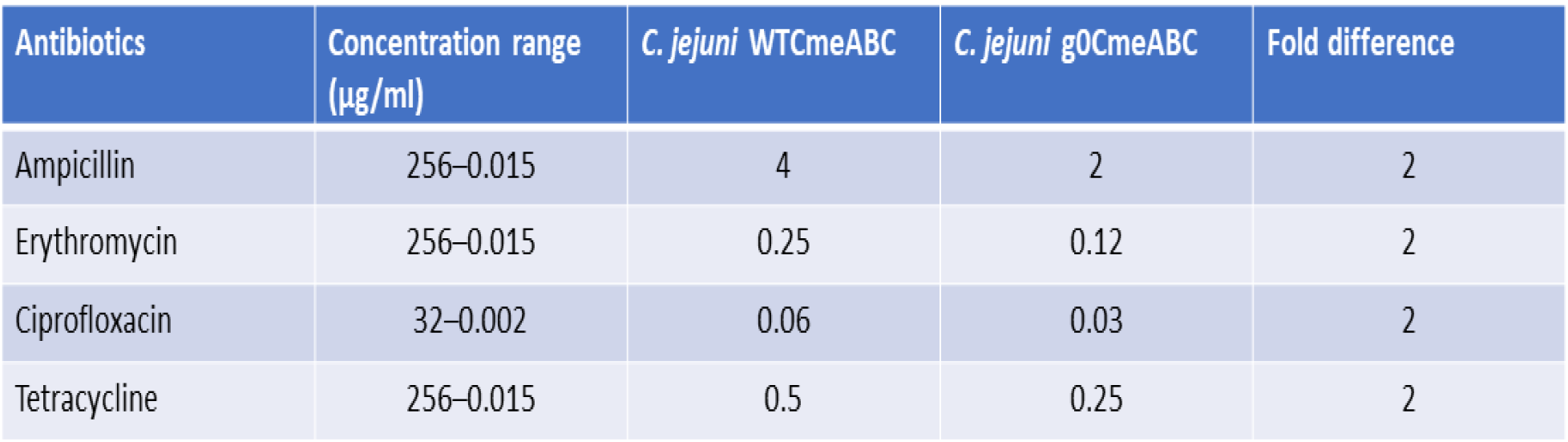
Minimum inhibitory concentration in C. jejuni strains. The minimum inhibitory concentration (MIC) of *C. jejuni* WTCmeABC and *C. jejuni* g0CmeABC was read directly from the strip at the point where the zone of inhibition of bacterial growth intersected with the antibiotic concentration on the strip. The results presented are the mean from three biological replicates two technical replicates each.

### Generation of fully glycosylated CmeA in glycocompetent *E. coli*

Previous studies showed that UDP-N-acetylglucosamine—undecaprenyl-phosphate *N*- acetylglucosaminephosphotransferase; WecA, could interfere with the biosynthesis of heterologous expression of polysaccharides built on the undecaprenyl-phosphate lipid anchor-rendering it built on incorrect glycan at the reducing end. To circumvent this problem and ascertain that g2CmeA is glycosylated with the native *C. jejuni N*-linked glycan we used glycocompetent *E. coli* SDB1^19^. The heterologous expression of an acceptor protein with protein glycosylation locus (*pgl*) usually yielded a mix population of glycosylated and non-glycosylated protein variant, indicating a suboptimal glycosylation process^20, 21^. We observed that *pglB* expression from pACYC(*pgl*) is insufficient to achieve optimal glycosylation (data not shown). To overcome this bottleneck, we sought boosting PglB expression by introducing pGXVN114 to *E. coli* SDB1 expressing CmeA and *N*-linked glycan biosynthetic pathway. Fig 2 Lane 1 shows the optimal glycosylation of constitutively expressed CmeA and *N*-linked glycosylation pathway along with IPTG inducible PglB from pGXVN114 backbone.

**Fig 2.**
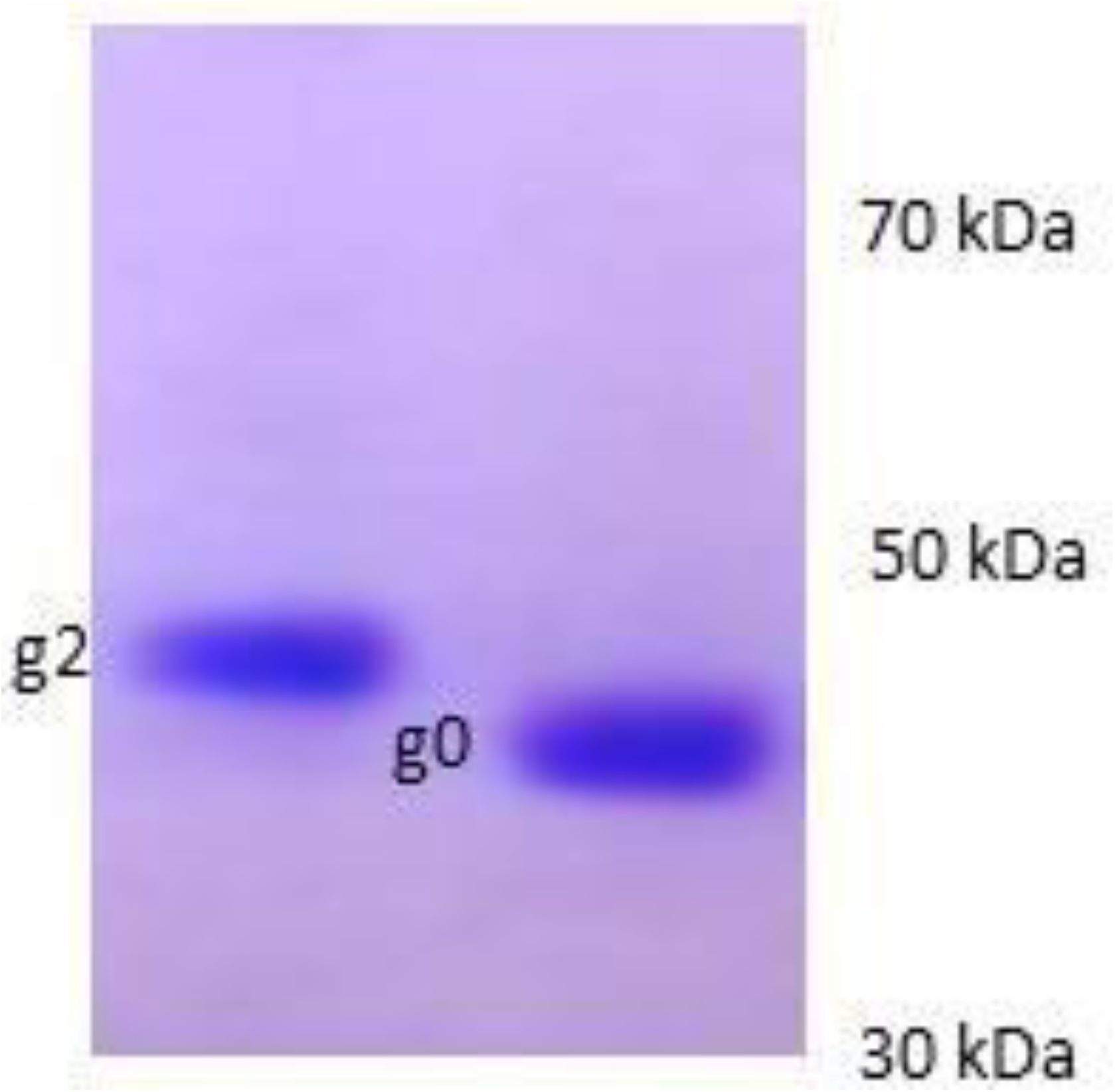
Generation of fully glycosylated CmeA in *E. coli* SDB1. CmeA variants were purified using IMAC followed by concentration and buffer exchange using Amicon ultra-0.5 ml centrifugal filter units. Proteins were then separated by SDS-PAGE and visualized by Coomassie blue staining.

### Glycosylation modulates protein global structure

In eukaryotes the enzymatic attachment of *N*-linked glycans to amide group of asparagine in the glycosylation consensus sequon, is correlated with changes in the biophysical properties of the protein such as thermostability^3^, aggregation, function^4^ and structure^22^. Glycans can confer interactions not only at the glycosylation site but also other regions in the protein. Serving as bulky hydrophilic groups, glycans can also favour certain conformational modifications that stabilise protein structure^22^. Bioinformatic studies suggested that glycans force the polypeptide chain to adopt a more extended conformation through restricting residual structures formation in the unfolded state^23, 24^. To investigate the role that *N*-linked glycans play in modulating the biophysical properties of CmeABC, we used circular dichroism (CD) spectroscopy. This allowed us to monitor the secondary structure as well as the conformational changes upon thermal denaturation of CmeA variants. Far-UV spectra for both g0CmeA and g2CmeA in 10 mM sodium phosphate, 75 mM sodium chloride, 10% glycerol buffer (pH 8) were collected at 20 °C. The spectrum exhibited helical structure signature with two negative minima at 208 and 222 nm. It also showed a positive maximum at 196 nm suggesting the presence of beta-sheets structure Fig 3 Superimposed CD spectra exhibit g0CmeA spectrum, shown in black, was slightly red shifted towards the β sheet. The CD spectra of both proteins were then analysed by BESTSEL^25^ for secondary structure content. Table 2 shows the secondary structure content of g0CmeA and g2CmeA at room temperature. Our results show that *N*-linked glycans confer subtle changes in protein global conformation.

**Fig 3.**
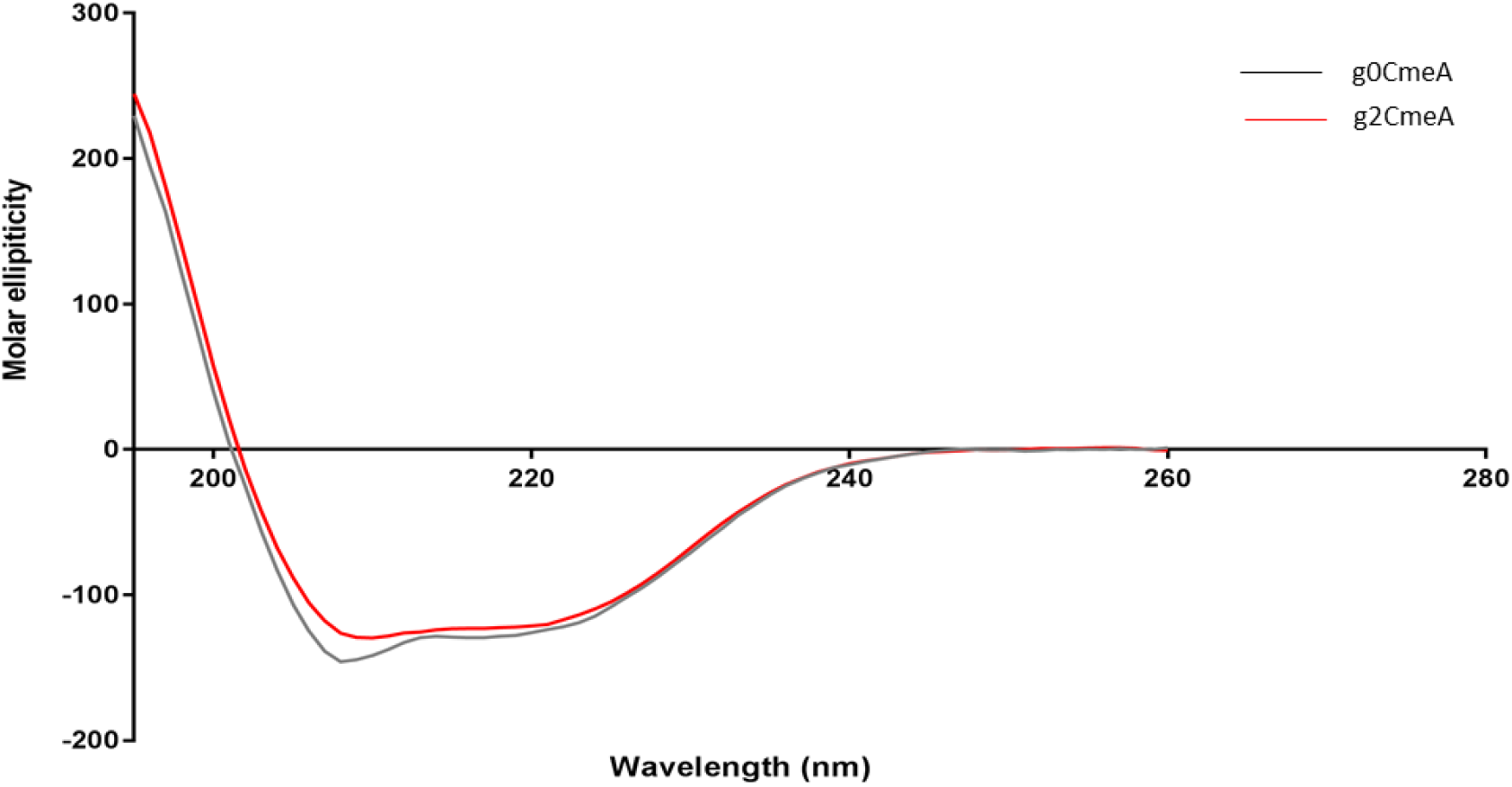
CD spectra of g0CmeA and g2CmeA in 10 mM sodium phosphate, 75 mM sodium chloride and 10% glycerol (pH 8.0). Far-UV CD spectra was collected for g0CmeA (0.124 mg/ml) and g2CmeA (0.174 mg/ml) variants in 0.5 mm rectangular cell path length. Molar ellipiticity was calculated and corrected for proteins concentration.

**Table 2.** Secondary structure calculation of g0CmeA and g2CmeA variants. CD units were converted to delta epsilon units and loaded to BESTSEL server. Although the conformations of both proteins are structurally similar, there is a subtle shift in the alpha helices and beta sheets ratios between both variants.

| | α-helix | β-sheets | Turn | Others |
| --- | --- | --- | --- | --- |
| g0CmeA | 29.6% | 26.4% | 11.9% | 32.1% |
| g2CmeA | 28.1% | 27.2% | 10.7% | 34.0% |

### Glycans enhances protein thermostability

It has been established that *N*-linked glycans play a role in stabilising glycoproteins thermodynamically in eukarotes^3, 22^. The intrinsic role of *N*-linked glycans in stabilising CmeA was investigated through analysing the CD spectra recorded for g0CmeA and g2CmeA at elevated temperatures. The multi–wavelength melting profiles monitored at 260- 195 nm were recorded during the heating of g0CmeA and g2CmeA from 6°C to 94°C at 1°C per minute rate with a 2°C step size. Isodichroic points were observed in the far-UV CD spectra at Fig 4, A **and** B supports more than two-state nature of the unfolding transition. Derivative of CD spectra were used to calculate the melting temperature (T_m_) of both g0CmeA and g2CmeA Suppl 1. The loss of CD spectra was observed upon incremental rises in temperature, melting curves measured for g0CmeA and g2CmeA show that both proteins have three transitional phases at T_m1_ 46.1°C ± 0.2, T_m2_ 53.5°C ± 0.4 and T_m3_ 56.7°C ± 0.6 for g0CmeA and T_m1_ 43.8°C ± 0.3, T_m2_ 49.3°C ± 0.2 and T_m3_ 62.8°C ± 0.2 for g2CmeA. No light scattering was observed in the UV spectra indicating turbidity presented at high temperature. This shift in final melting temperature suggests that glycans stabilise g2CmeA at elevated temperature Fig 4, C.

**Fig 4.**
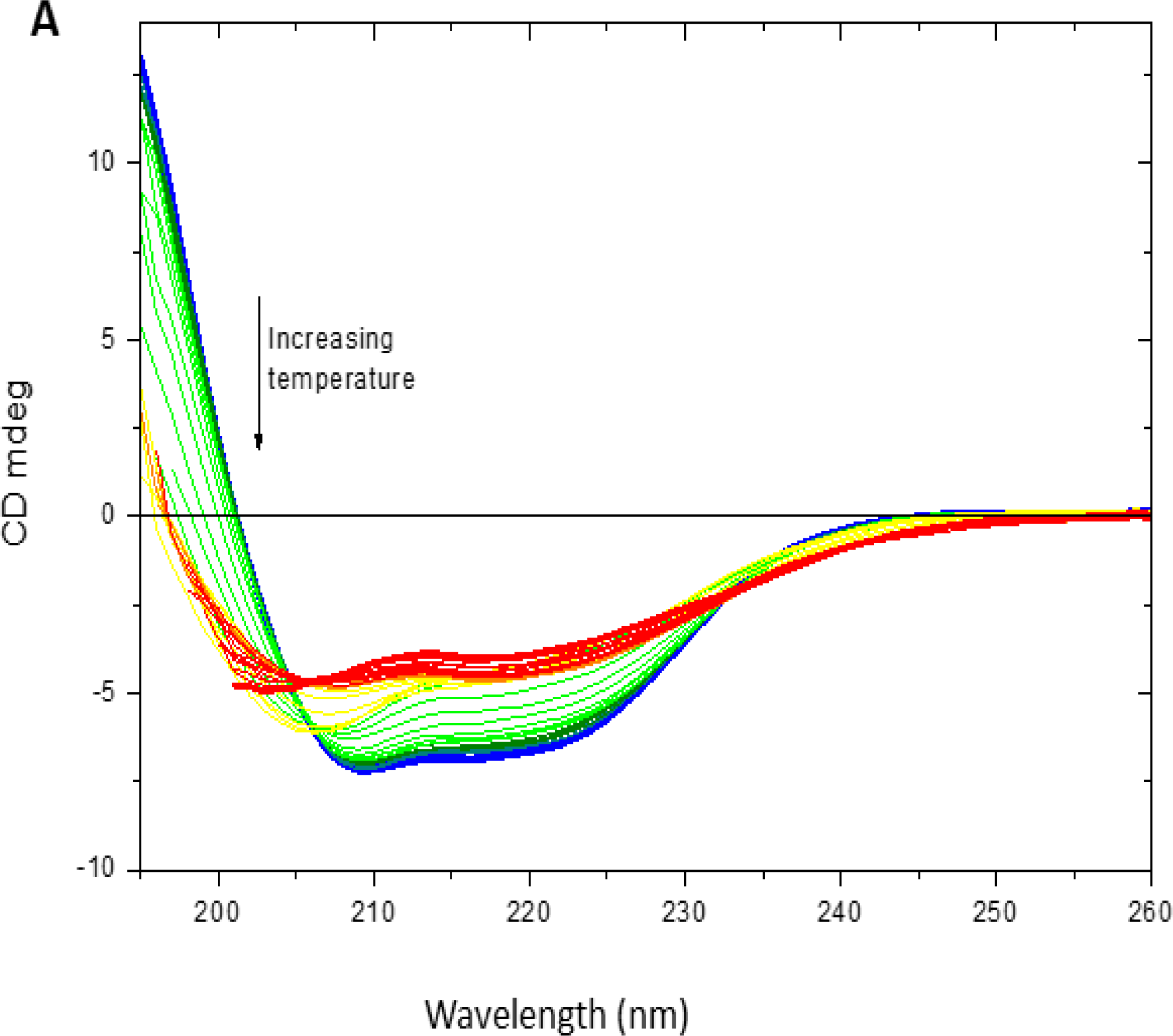
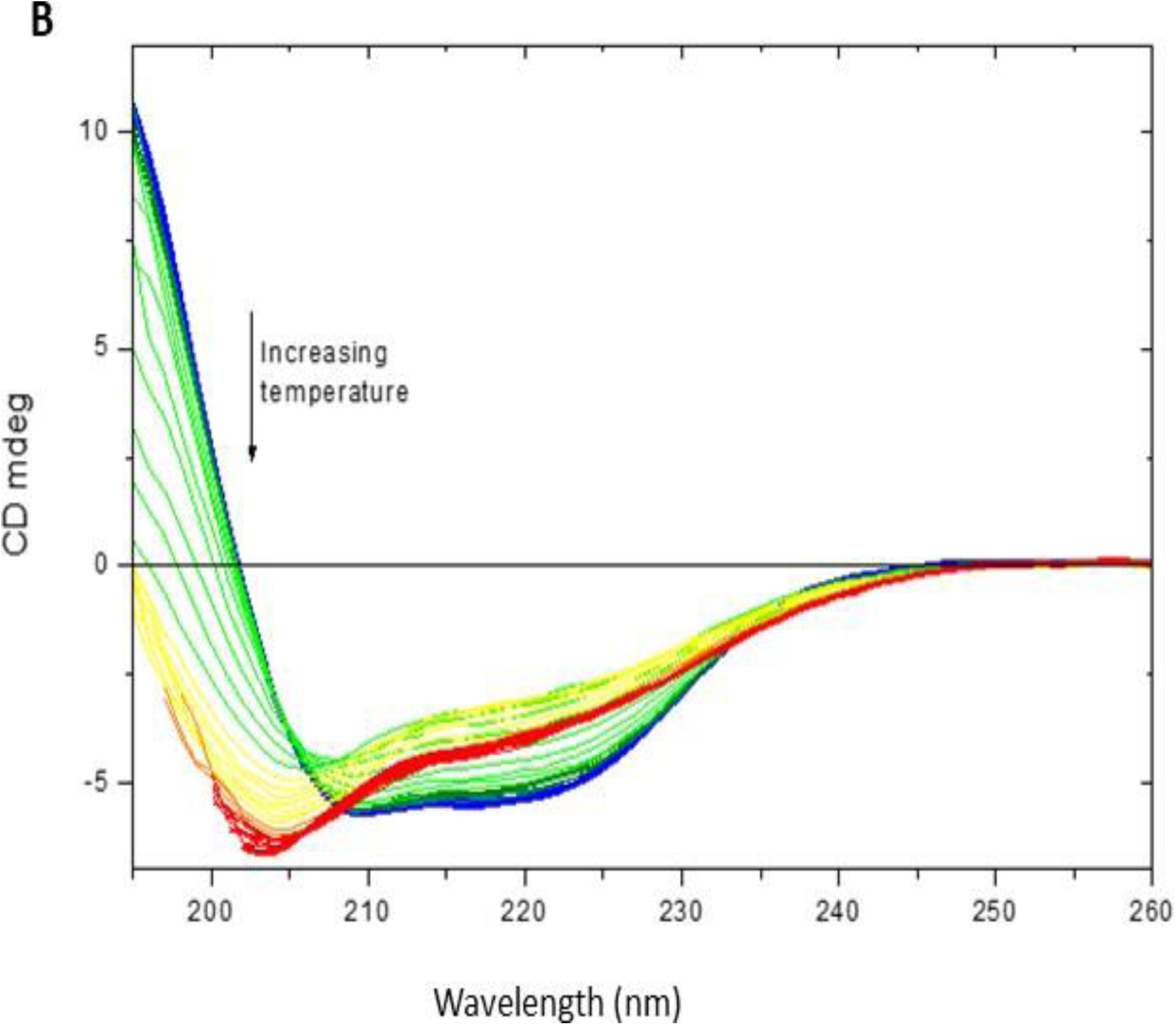
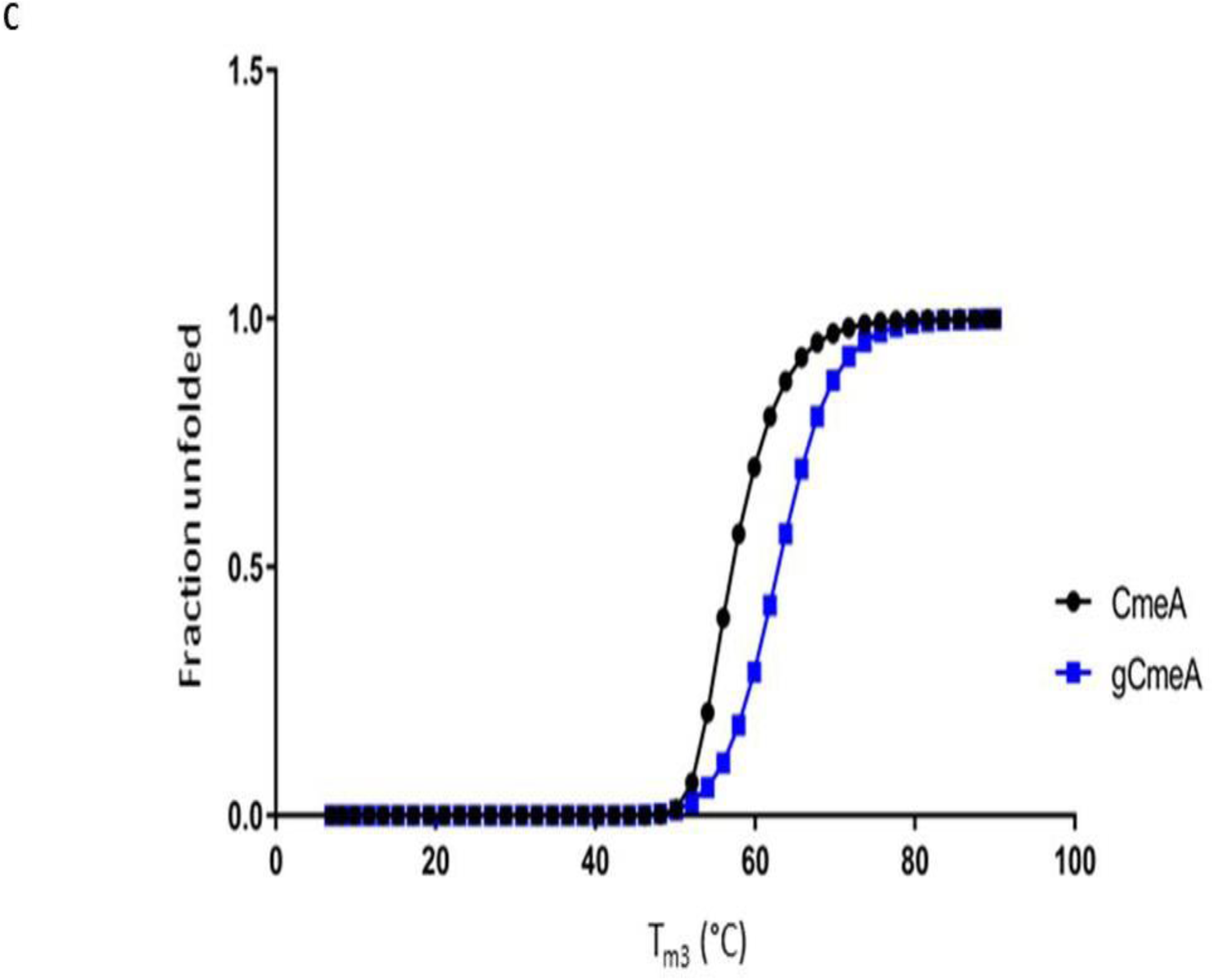
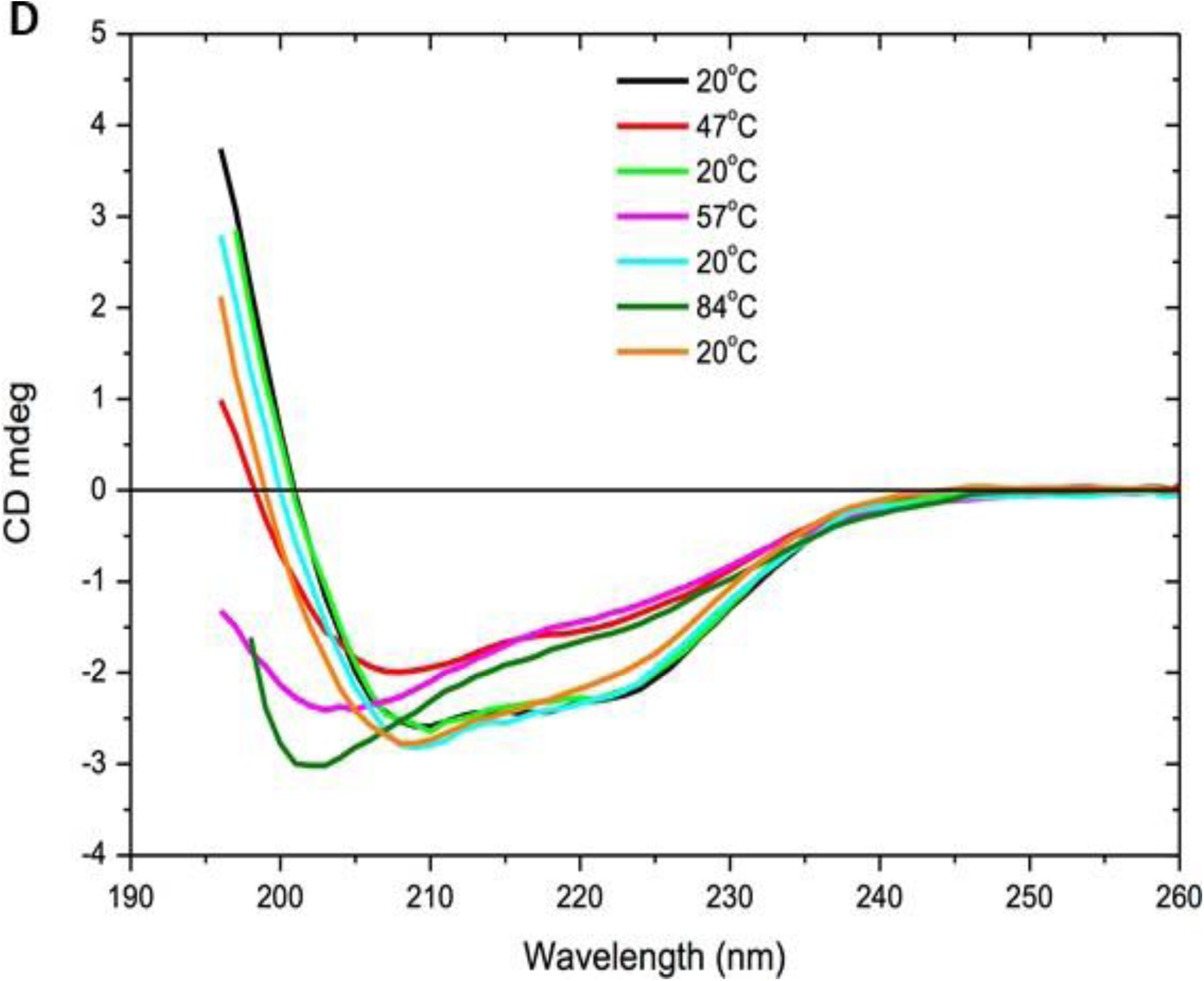
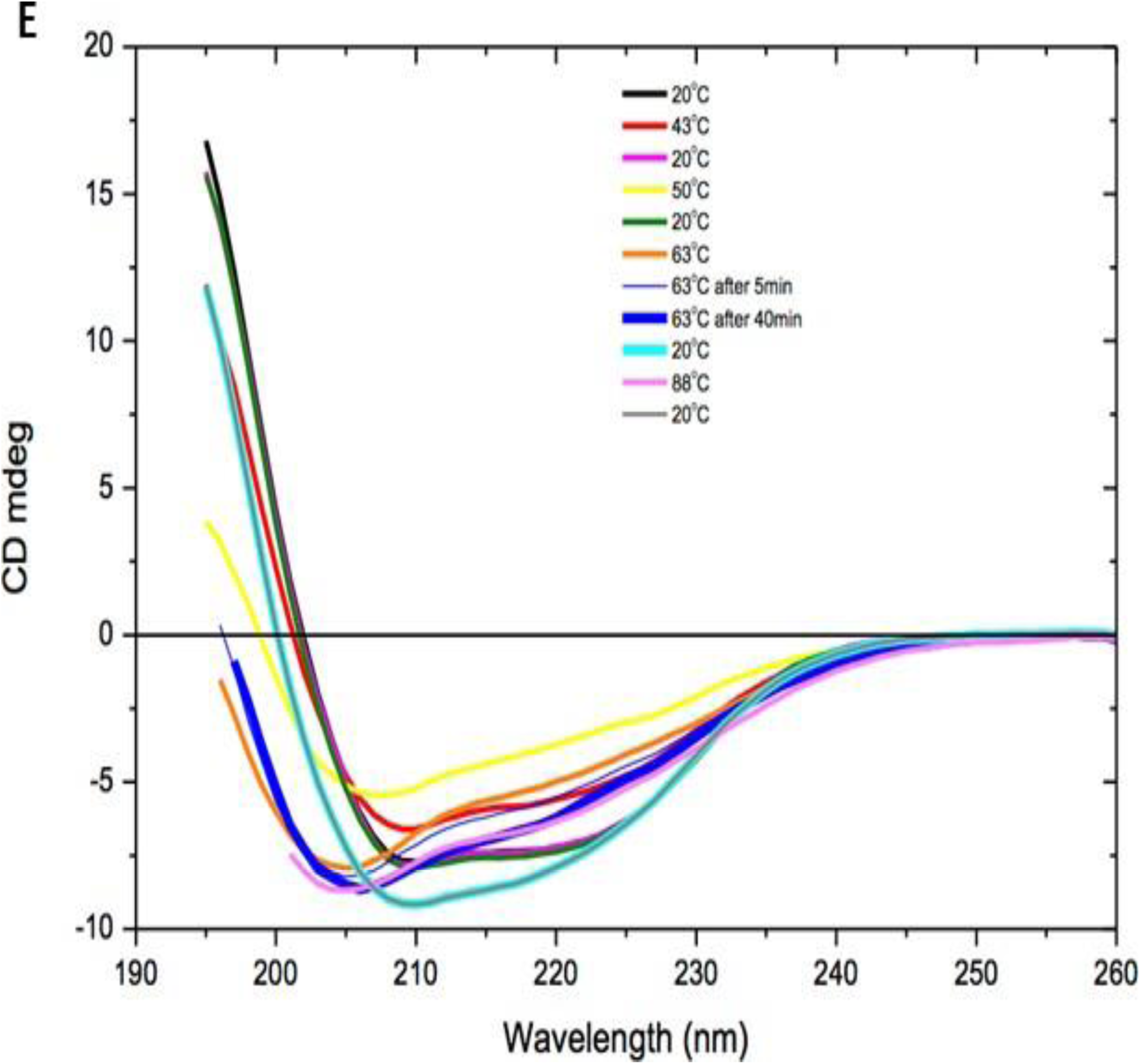
Thermal melts of g0CmeA and g2CmeA in 10 mM sodium phosphate, 75 mM sodium chloride and 10% glycerol (pH 8.0). Far-UV CD spectra was collected for g0CmeA (0.124 mg/ml) and g2CmeA (0.174 mg/ml) variants in 0.5 mm rectangular cell path length. CD mdeg were recorded as a function of temperature from blue (6 °C) to red (94 °C) for g0CmeA, (A) and g2CmeA (B). Each colour in between was obtained at rate 1 °C per minute with a 2 °C stepwise increase.CD spectra to asses the reversibility of thermal unfolding study was recorded at 20°C, raised to T_m_ and re-cooled to 20°C sequentially. CD spectra was collected for 5 minutes at each temperature interval for g0CmeA, (C); Thermal denaturation of CmeA (black) and g2CmeA (blue) as change of ellipticity values upon temperature increase (D); g2CmeA (E). CD spectra of g2CmeA was stabilised after 30 minutes at T_m3_ indicating a more resilient behaviour thermal unfolding process.

To confirm the previous findings, we examined conformational folding reversibility and unfolding rate for both g0CmeA and g2CmeA. The assay is based on three successive cycles whereby CmeA variants were cooled at 20°C, heated up to the corresponding T_m_ for 5 minutes and then cooled again at 20°C. To assess conformational folding reversibility, CD spectra that were recorded at 20°C, before and after increasing the temperature to the corresponding T_m_ were compared. CD spectra of CmeA variants were superimposable before and after the first two cycles of heating (T_m1_ and T_m2_) but not after heating at T_m3_, indicating conformational changes Fig 4, D **and** E. Unfolding rate was evaluated according to changes in CD spectra with respect to time at T_m3_. A significant reduction in CD spectra was observed when g0CmeA and g2CmeA were heated at their corresponding T_m3_. The unfolding of CmeA was achieved in 5 minutes at its T_m3_. Notably, the CD spectra recorded for g2CmeA at its corresponding T_m3_ kept changing for 30 minutes indicating a slower unfolding rate. This result along with the above data highly suggest that *N*-linked glycans play a pivotal intrinsic role in protein thermodynamic stabilisation.

### Glycans modulate molecular assembly and protein-protein interaction

Unlike eukaryotes, there is no evidence that *N*-linked glycans modulate protein-protein interaction or complex assembly in prokaryotes. To explore the potential role of *C. jejuni* general *N*-linked glycans in modulating the interaction of glycoproteins with their cognate partners, surface plasmon resonance (SPR) was used. SPR was previously used to investigate the interactions in orthologous membrane fusion proteins (MFP) with TolC from of *E. coli*^26^. Quantitative analysis of the binding curves showed multiple reaction events. The model used to determine binding kinetics indicated the presence of two populations of MFP proteins. The two populations exhibited different binding kinetics, notably, fast and slow dissociation rates that contributed to weak and strong interactions, respectively^26^. We employed a CM5 chip with g0CmeA and g2CmeA immobilised through amine coupling, CmeC was then injected over CmeA variants surfaces in different concentrations. In our model, g0CmeA and g2CmeA exhibited multiple events interaction with CmeC. These interaction events can be attributed to a fast and a slow association and dissociation rates. Quantitative analysis of the sensogram yielded excellent results for slow interactions however, fast interactions could not be fitted in a model to generate accurate binding kinetics. At pH 7.4, both CmeA variants exhibited similar dissociation rate constants (k_off_) of 9e^−4^ s^−1^ for g0CmeA and 7.5e^−4^ s^−1^ for g2CmeA Fig 5, A **and** B. Interestingly, difference in association rate constant (k_on_) was observed, g0CmeA k_on_ = 5e^4^ (M^−1^s^−1^) whilst g2CmeA k_on_ =1.5e^5^ (M^−1^s^−1^). This difference in the k_on_ rate indicates that g2CmeA possess more binding pockets that allows slow yet high affinity interactions with CmeC compared to g0CmeA. The equilibrium dissociation rate constants (K_D_) derived from the binding kinetics analysis were 1.7e^−8^ (M) and 5e^− 9^ (M) and g0CmeA and g2CmeA, respectively.

**Fig 5.**
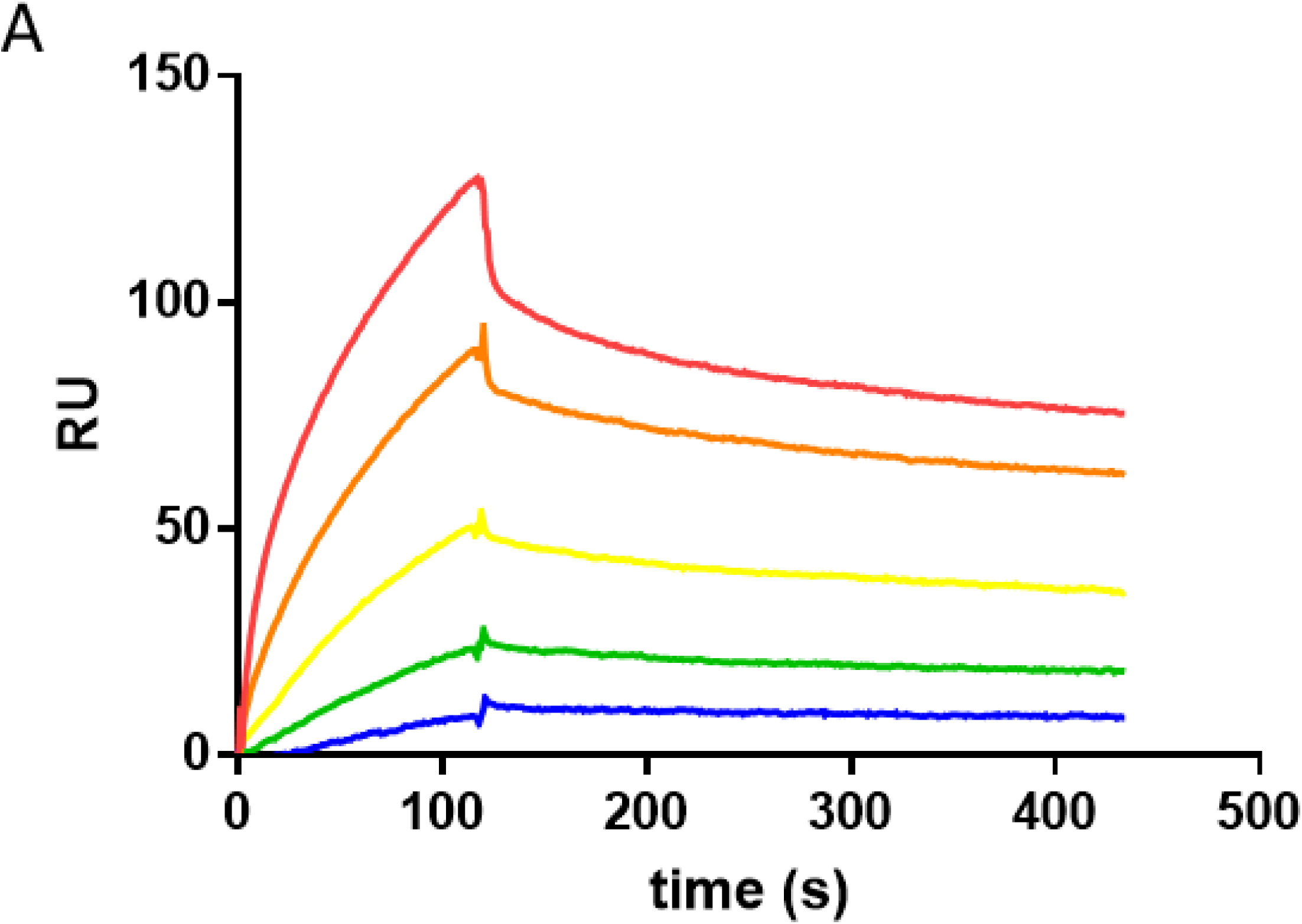
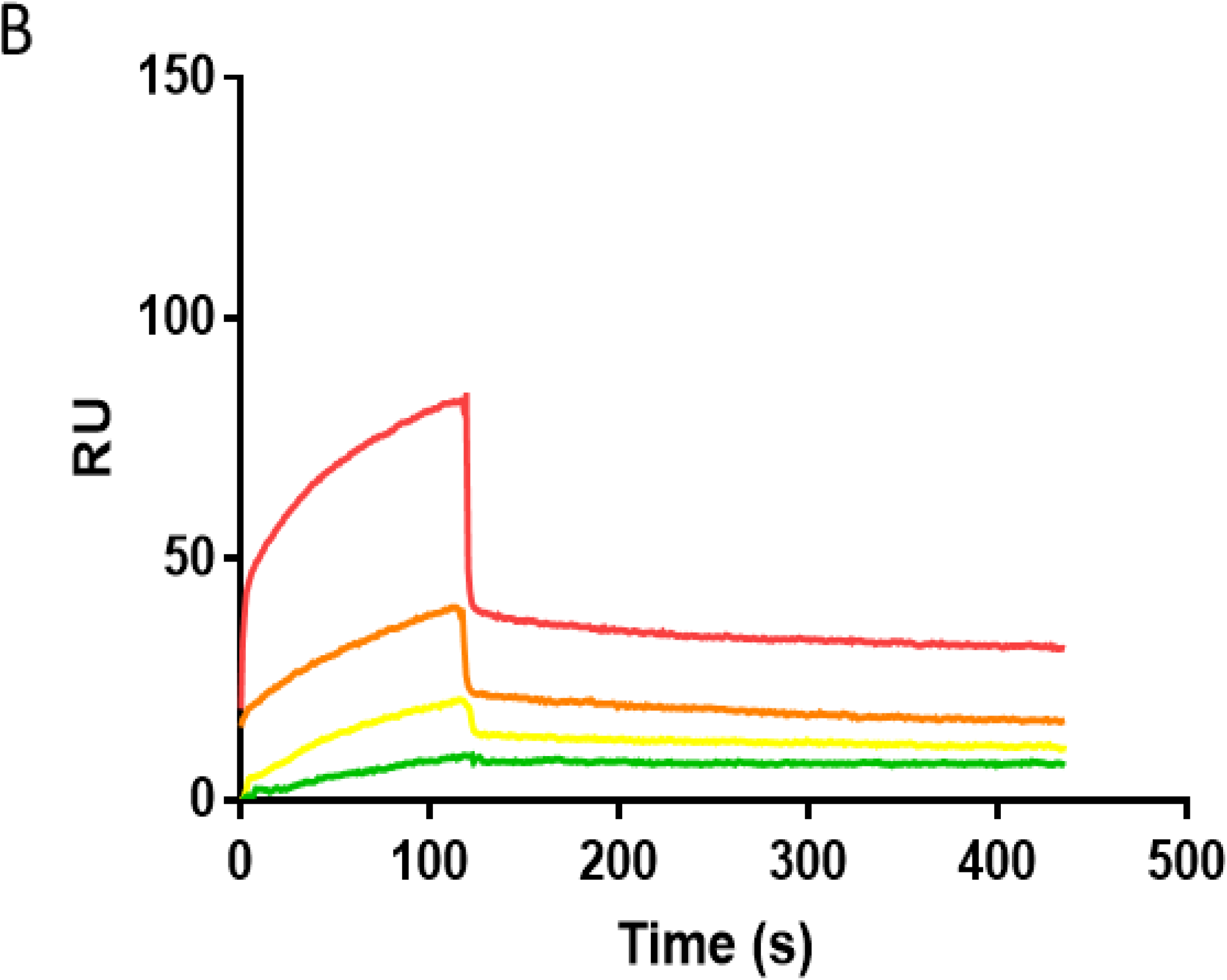
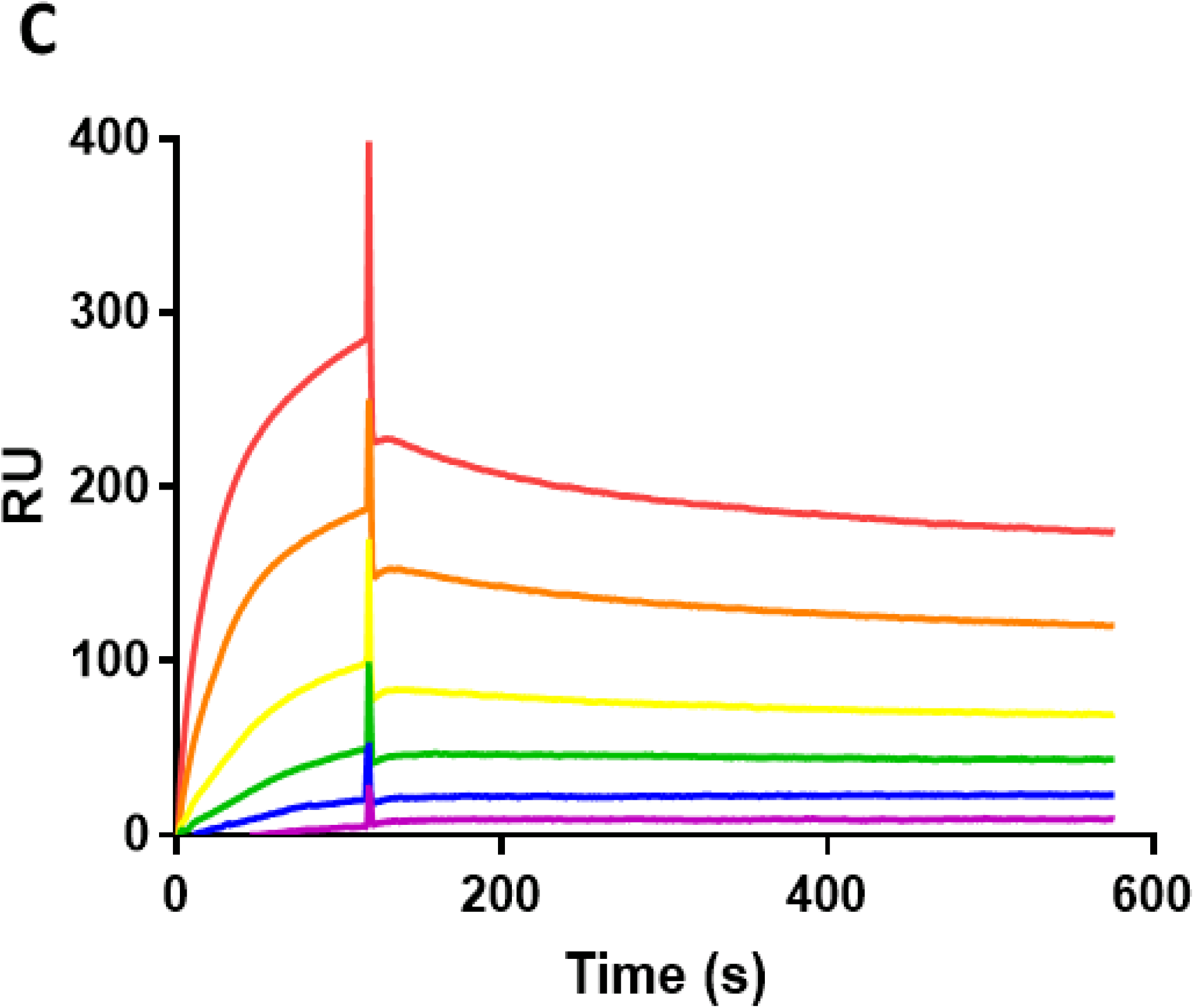
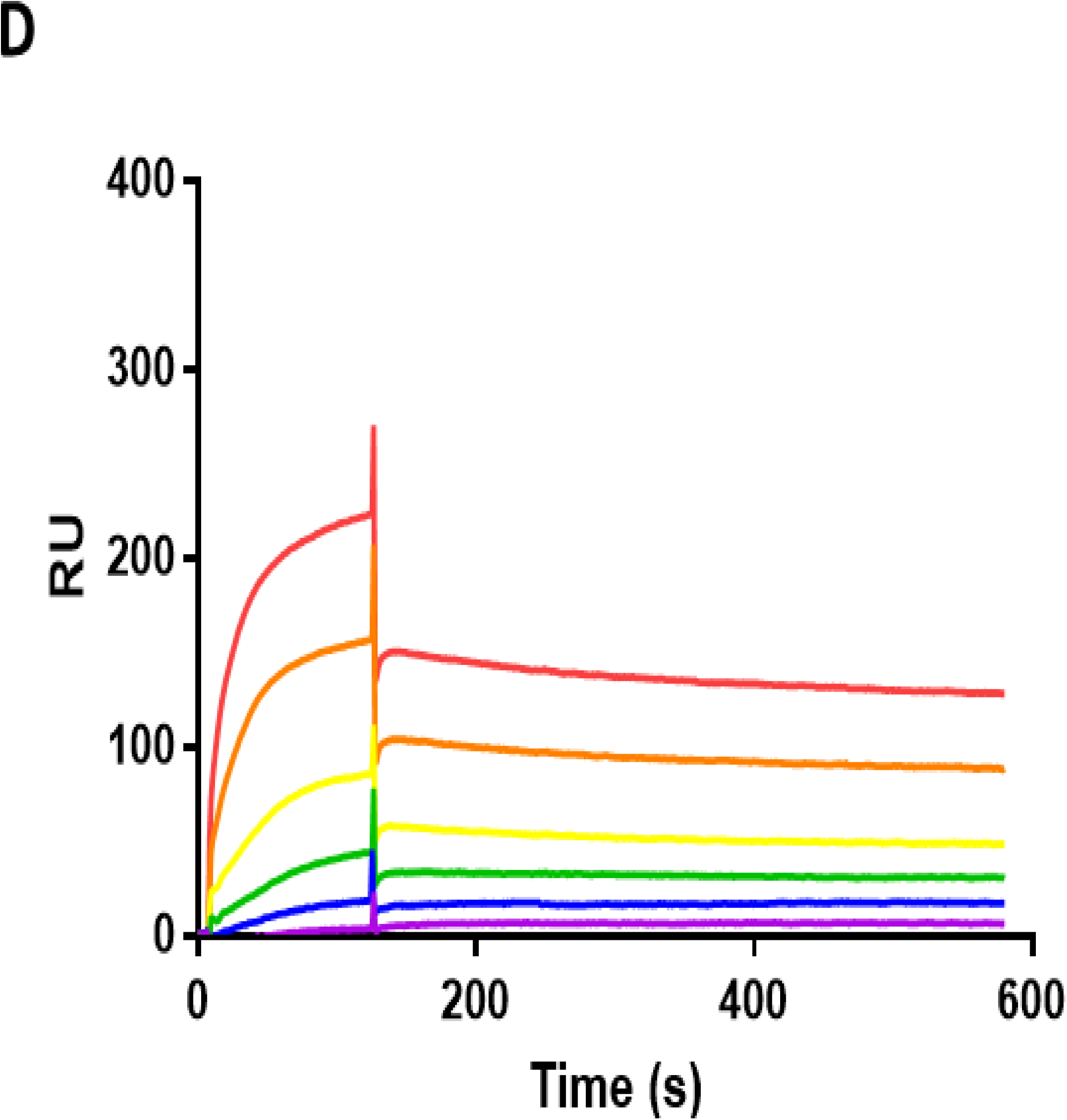
Glycosylation enhances interactions between CmeA variants and CmeC. SPR analysis of CM5 chip with A) 900 RU of g2cmeA immobilised and B) 1040 RU of g0cmeA immobilised. Association of CmeC at pH 7.4 was performed for 2 mins and dissociation was followed for 5 mins. Concentrations of CmeC were two-fold dilutions from 2×10^−7^ M (red) to 1.25 ×10^−8^M (blue) or 2.5 ×10^−8^ M (green). SPR analysis of CM5 chip at pH 6.0 with C) 900 RU of g2CmeA immobilised and D) 1040 RU of g0CmeA immobilised. Association of CmeC was performed for 2 mins and dissociation was followed for 5 mins. Concentrations of CmeC were two-fold dilutions from 2−10^−7^ M (red) to 0.6 ×10^−8^M (purple)

To investigate the effect of pH on modulating binding kinetics, we observed CmeA- CmeC interactions at pH=6.0 Fig 5, C **and** D. At this pH CmeA-CmeC interactions were more avid and with a greater number of sites bound. Similar to binding curves observed at pH 7.4, g2CmeA showed a favourable slow association and dissociation binding curves than g0CmeA. The number of sites for slow interaction were greater for g2CmeA contributing to a modestly higher affinity for interaction with CmeC compared to a weaker affinity for CmeC exhibited by g0CmeA. To confirm that variations in binding kinetics were not due to differences in structural orientation between g0CmeA and g2CmeA, both proteins were immobilised on NTA chip using C-terminal 6xHis tag, CmeC was then passed in different concentrations. Binding kinetics indicated similar k_on_ and k_off_ for both CmeA variants, although fewer sites were available Suppl 2. Interestingly, g2CmeA bound more CmeC than g0CmeA, confirming the data seen with amine coupling. These results show a complex binding pattern between CmeA variants and CmeC. They also suggest an extrinsic role for *N*- linked glycans, exhibited in the variation in binding kinetics between g0CmeA and g2CmeA, where the glycosylated form of CmeA showed a greater proportion of higher affinity interaction sites than its non-glycosylated counterpart.

## Discussion

Whilst the role of glycosylation in eukaryotes has been thoroughly explored, a similar depth of investigation is lacking in prokaryotes. The presence of different glycosylation systems in prokaryotes has been regularly reported^2, 7, 11–13^. Some have described their effect in virulence, adhesion and motility by creating genetic knock outs of the glycosylation machinery^10–12, 32, 33^. Nonetheless, these reports have not provided in-depth studies into the direct role that *N*-linked glycans exert on protein function. Glycoproteomic studies revealed diBacNAc to be conserved across *Campylobacter* species. Notably, diBacNAc was also found to be at the reducing end of *O*-linked glycans in *Neisseria gonorrhoeae*, indicating a parallel evolution between *N*-linked and *O*-linked glycosylation systems in bacteria^8, 9, 34^. The role of *N*- linked glycans in stabilising major multidrug efflux pump in *C. jejuni* has been shown to contribute in efficiently extruding antimicrobials and ethidium bromide. Disrupting glycosylation in CmeABC resulted in higher accumulation of ethidium bromide and lowering antibiotic MIC in *C. jejuni*. These differences in activity are not due to the low abundance of CmeABC complex in g0CmeABC strain Fig 1, A. A protein synthesis arrest assay showed that loss of glycosylation did not promote CmeC proteolytic degradation. This is in agreement with the previous finding that CmeC protein abundance were equal in *C. jejuni* and *C. jejuni pglB*::*aphA* (Abouelhadid S, *et al* paper submitted). This result suggests that *N*-linked glycans might be modulating molecular assembly. Studies on truncated *N*-linked glycans will reveal the role of each glycostructure, it will also help to understand the role of the conserved first two glycans between different *Campylobacter* strains.

Bioinformatic studies investigating protein structural changes exerted by glycans has been inconclusive. These studies rely heavily on the *in-silico* analysis of protein structure entries in the protein database bank. Whilst modern advances in crystallographic techniques pave the way for more structural studies, obtaining glycoprotein structure is still challenging and remains poorly represented in the protein database bank. Xin *et al* reported that protein glycosylation causes significant yet unexpectedly subtle changes in both local and global protein structure (up to 7%) ^23^. However, Hui Sun lee *et al* concluded that *N*-glycosylation causes non-significant changes in protein structure but increases protein stability likely due it a role played in reduction of protein dynamics^24^. Experimentally, our initial CD study of CmeA variants showed that both have the same conformational fold, however they confer subtle structural differences to the protein Fig 3, B. A small shift has been observed in the percentage of alpha helices and beta sheets between g0CmeA and g2CmeA, 1.3 ±0.2. It is still unclear whether the structural variations are due to local stabilisation resulting from the glycosidic bond between the asparagine side chain in the glycosylation site and *N*-linked glycans, or global structural rearrangement due to the interaction of the glycan with other distant regions in the protein backbone. A structural elucidation of CmeA in its glycosylated and non-glycosylated might provide insights on the extent of the importance of conformational changes.

It has been suggested that *N*-linked glycans might enhance protein thermostability. The glycoprotein PEB3 from *C. jejuni* was used to test the stabilisation effect of *N*-linked glycans. Average melting temperature of PEB3 (K135E) variants were analysed using SYPRO orange thermoflour. Interestingly, the T_m_ of glycosylated PEB3 was shown to be 4.7°C higher than its non-glycosylated counterpart; PEB3^35^. This comes with an agreement with CD thermal melts of g0CmeA and g2CmeA. CD thermal melts showed that whilst both of the CmeA variants have the same apparent unfolding behaviour, T_m_ of g2CmeA was 6.4°C ±0.5 higher than that of g0CmeA. The three transitional phases of both variants showed that g2CmeA seems to be responding to a rise in temperature via conformational rearrangements at 2.4°C ±0.1 lower than g0CmeA Fig 4, A. CD spectra recorded after cooling showed that the structural rearrangements were reversible and the protein could fold again, suggesting that protein fold/unfolding T_m1_ and T_m2_ are reversible for both of the CmeA variants. Remarkably, the unfolding behaviour of g2CmeA at T_m3_ was different to g0CmeA at its correspondent T_m3_ in the conformational reversibility assay. Time taken to unfold g2CmeA was at least 5 times more than g0CmeA, thus indicating a role played by *N*-linked glycans in conferring greater resistance to unfolding Fig 4, C **and** D. We postulate that *N*-linked glycans stabilise g2CmeA through a reduction in the unfolding rate in g2CmeA, this finding agrees with the observation that eukaryotic *N*- linked glycans stabilise the hCD2ad through slowing the unfolding rate of the protein 50- fold when compared to its non-glycosylated counterpart^2^.

Owing to the lack of subcellular compartments, the extrinsic role of prokaryotic *N*- linked glycans in protein-protein interaction has been not fully appreciated. Despite the scarcity of glycoproteomic data, few molecular assemblies have been reported to have at least one of its component to be glycosylated ^17, 31^. We demonstrate a potential extrinsic role of *N*- linked glycans in CmeA interaction with CmeC. In an orthologous multidrug efflux pump; AcrAB-TolC, AcrA showed the presence of two populations, of the same protein, interacting with different kinetics to TolC. The two populations contributed to a fast weak interaction and slow strong interaction Fig 5, A **and** B. The complexity of these interactions is exaggerated in *C. jejuni* due to the presence of *N*-linked glycans, that could modulate interaction of CmeA with CmeC. Quantitative analysis for the interaction kinetics of CmeA variants with CmeC showed that, *N*-linked glycans increase the binding affinity to CmeC by 3.4-fold. That was clearly demonstrated in the difference in K_D_ between CmeA variants at pH= 7.4. The difference in binding affinity was confirmed when CmeA variants were immobilised with the same orientation on Ni chip. Recently, a pseudoatomic structure provided a detailed picture of interaction between AcrA and TolC. This elaborated the adaptor bridging-binding model that involved an intermesh cogwheel-like binding between AcrA and TolC^36^. The conserved binding motif Val-Gly-Leu/Thr (VGL) is located at the tip region of the coiled coil α-hairpin of the protein, serving as a site of interaction with the RXXXLXXXXXXS (RLS) motif of AcrA^36^. In light of this study, our computational analysis showed that CmeC from *Campylobacter spp* does contain a truncated VGL motif, denoted VGA, whilst we found the RLS motif to be conserved in among *C. jejuni, C. lari, C. coli* and *C. fetus* Fig 6, A. To understand whether *N*-linked glycans modulate protein-protein interaction we analysed the proximity of glycosylation sites to VGA and RLS binding site in both CmeC and CmeA, respectively. The glycosylation sites in CmeC were shown to be distant from the proposed binding site Fig 6, C and probably closer to the transmembrane domain of the protein. Remarkably, we found that one of the glycan modified asparagine (^123^N) is at the X_−1_ position to RLS motif and is conserved in *C. jejuni* and *C. coli*, but not in *C. lari* and *C. fetus* Fig 6, A **and** C. This strongly suggests that the localisation of *N*-linked glycan adjacent to RLS might be affecting either the local site conformation and/or promote a stronger interaction with the VGA motif in CmeC; resulting in the interaction kinetics differences between g2CmeA and g0CmeA with CmeC observed by SPR in this study. In eukaryotes, it is established that *N*-linked glycans at different glycosylation sites in the same protein could play different roles. The roles of these *N*-linked glycans can be categorised into; (a) promoting protein folding, (b) modulating protein trafficking and localisation and (c) effecting protein functionality.

**Fig 6.**
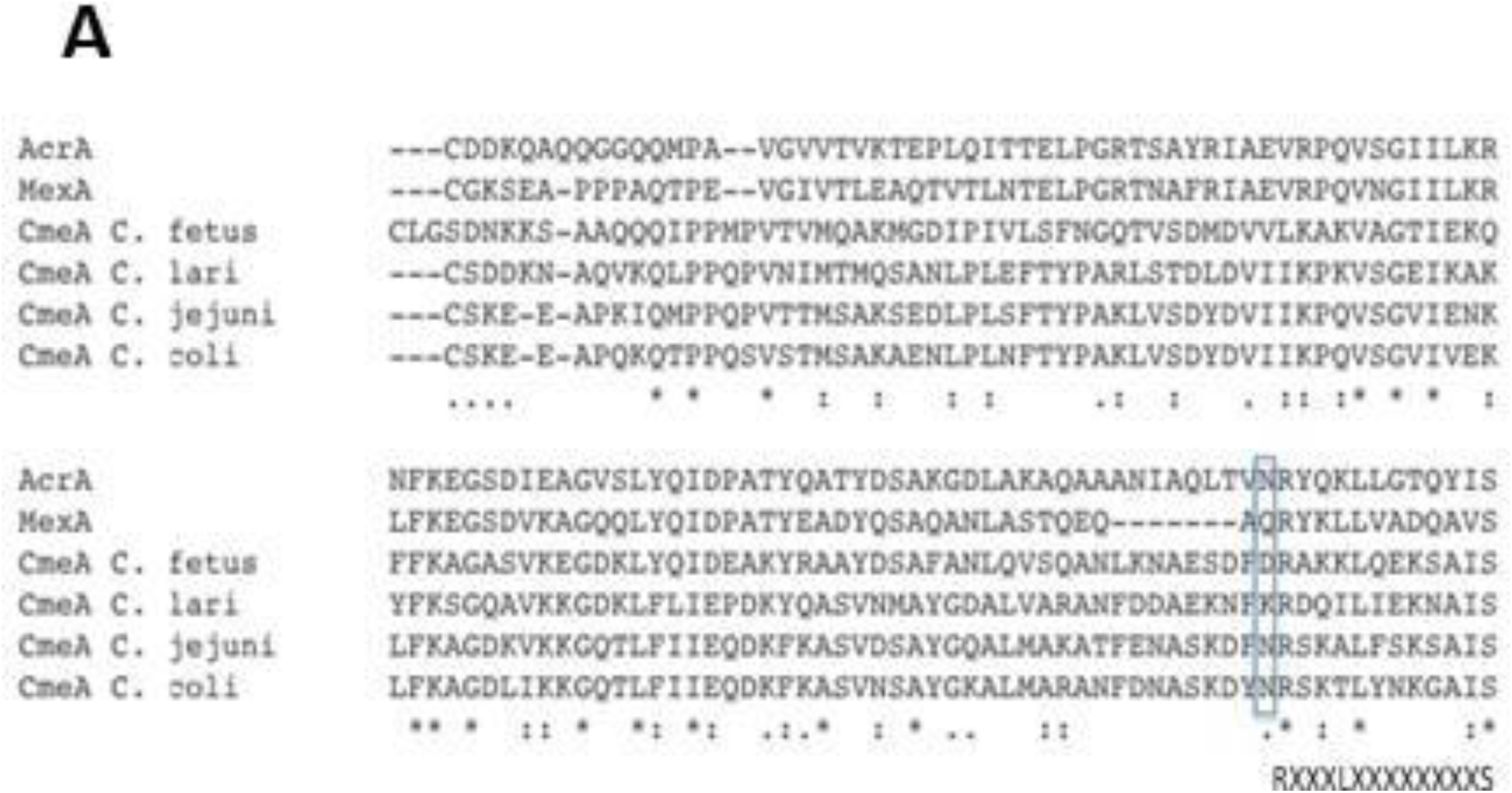
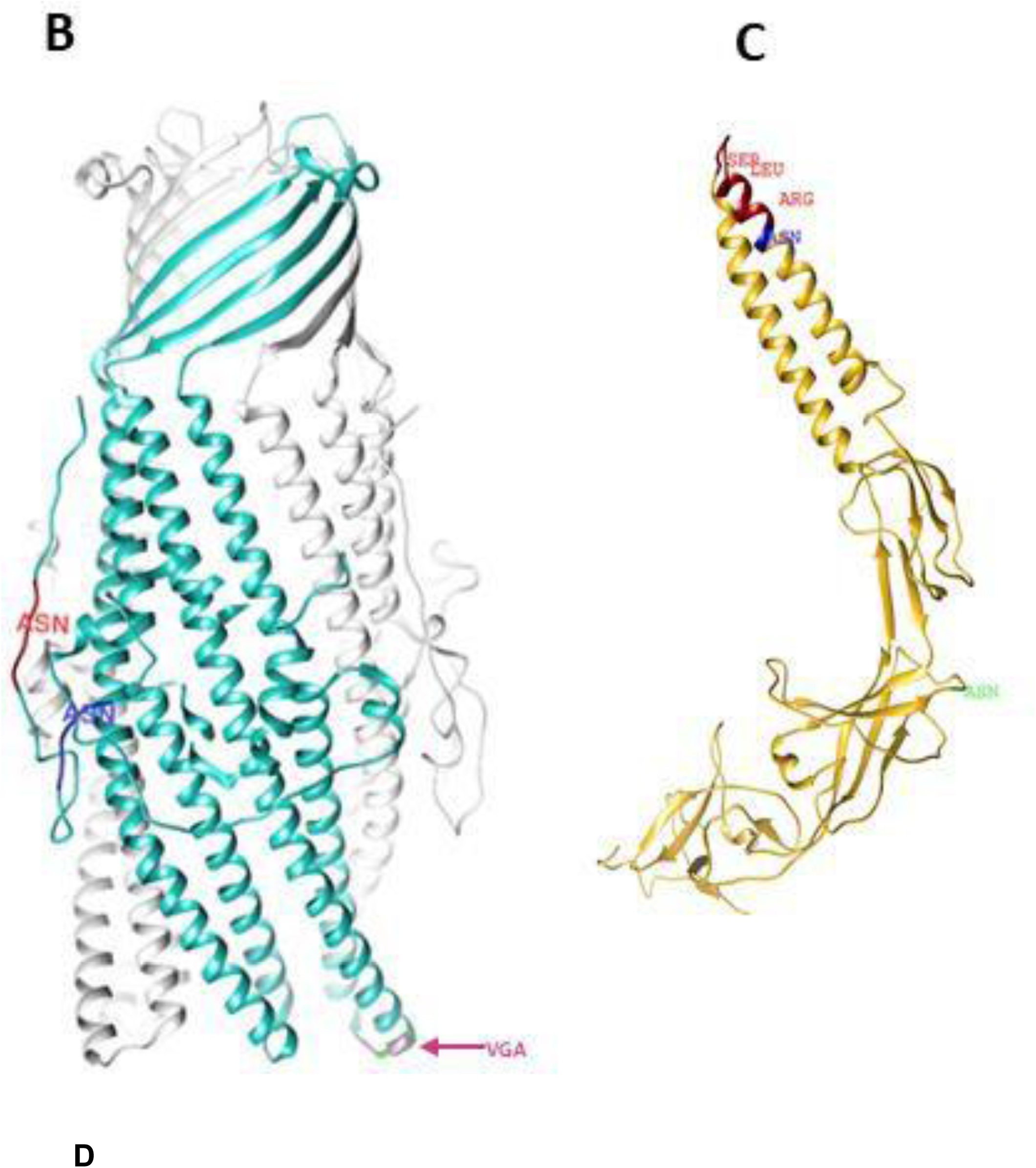
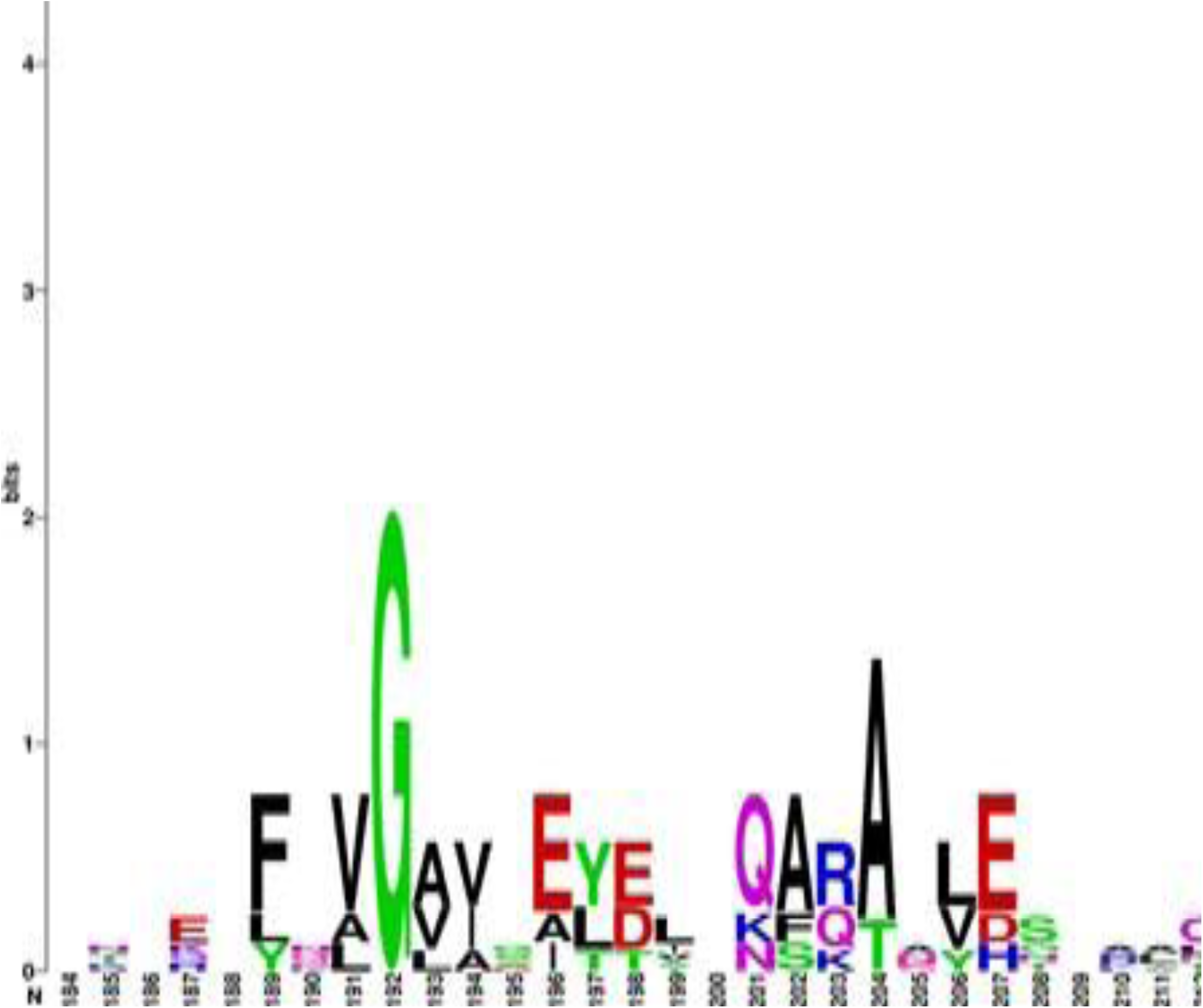
Analysis of binding sites in CmeA and CmeC (A) Amino acid alignment of signal peptide processed CmeA orthologues. Conserved amino acids are denoted by an asterisk, similar amino acids are denoted by colon and weak amino acid similarity is denoted by period. The amino acid sequences were retrieved from Uniprot and aligned using Clustal Omega^42^. RLS attachment site is shown to be conserved among periplasmic accessory proteins from different strains. The localisation of N is highlighted in blue box, showing the presence of ^123^N at X_−1_ to the conserved RLS motif in *C. jejuni* and *C. coli* but not *C. fetus* nor *C. lari*. (B) Structural representation focusing on chain A of CmeC trimer (PDB:4MT4). Chain A is highlighted in cyan, ^32^N and ^49^N are highlighted in red and blue respectively. The proposed attachment site VGA motif is highlighted in magenta showing its distant from both of the glycosylation sites. (C) Structural prediction of CmeA. Signal processed amino acid sequence was deposited in I-TASSER and the best structural fit was based on MexA model(ref). RLS motif is highlighted in dark red, ^123^N and ^273^N are highlighted in blue and light green, respectively showing the close proximity of ^123^N to RLS motif in CmeA. (D) Analysis of outer membrane channel; CmeC, AcrA and OrpM showing the conservation of Gly structurally located at the tip region of coiled-coil α hairpin domain among *Campylobacter* species, *E. coli* and *P. aeruginosa*.

This study provides the first detailed analysis of the role of bacterial *N*-linked glycan. The role of bacterial general *N*-linked glycans has been difficult to elucidate. This led to a notion that bacterial general *N*-linked glycans do not play any role in protein folding or function^39^. This notion was based on previous inconclusive results on the role of bacterial general *N*-linked glycans in modulating proteins function^37–40^. Our work refutes this widely held notion and demonstrates that bacterial *N*-linked glycans do not only play a role in slowing protein unfolding process and enhancing its thermostability but also it modulates protein interaction with its cognate partner. It also demonstrates a conserved role of general *N*- linked glycans previously seen in eukaryotes. This also suggests a common evolutionary role that led to the emergence of *N*-linked protein post translation modification, in expanding the functionality of proteome repertoire across all domains of life. Our proposed model can be used to interrogate prokaryotic general. glycosylation systems Our proposed model can be used to interrogate prokaryotic general glycosylation systems and help in the development of novel antimicrobials.

## Materials and methods

### Bacterial strains and growth conditions

*Campylobacter jejuni* 11168^10^ and its derivatives; *C. jejuni cmeD*::*cat*, *C. jejuni cmeD*::*cat wtcmeABC* and *C. jejuni cmeD*::*cat* g0*cmeABC* were used in this study. *C. jejuni* 11168H was grown on Columbia based agar or Muller Hinton based agar supplemented with 5% horse blood according to manufacturer’s instructions. Strains were grown at 37°C in a variable atmospheric incubator (VAIN) cabinet (Don Whitely, UK) maintaining microaerophilic conditions of: 85% Nitrogen, 5% Oxygen and 10% carbon dioxide. All of the cloning experiments were done in *Escherichia coli* DH10Beta (New England Biolabs, USA). *E. coli* DH10B was used in expression of CmeA and cloning and expression of CmeC whilst gCmeA was expressed in *E. coli* SBD1. *E. coli* strains were grown on either Luria-Bertani Broth or Luria-Bertani Agar and antibiotics were added when necessary.

### Inactivation of *cmeD* and generation of *C. jejuni cmeD::cat*, *C. jejuni cmeD::cat wtcmeABC* and *C. jejuni cmeD::cat g0cmeABC*

The nucleotide sequence of *cmeD* gene was commercially synthesized (Clonetech, USA) to also carry a chloramphenicol resistant gene; *cat* was inserted in the middle of *cmeD* to disrupt the gene. The DNA was then released by restriction digestion with EcoRV and cloned in pJET1.2 -following manufacturer’s instructions- to give pATN. Cloning of *cmeABC-aphA* was achieved by the following; *cmeABC* locus was amplified by primer FWDCmeA and primerREVCmeC with Phusion polymerase (New England biolabs, UK) using *C. jejuni* 11168H genomic DNA as a template, 6Xhis tag was added at the C-terminus of the CmeC to track its expression. The PCR amplicon was cloned in pJET1.2 following the manufacture’s instructions to give pMH3 that was then cut by BamHI to introduce the kanamycin resistant gene *aphA*, to be used as an antibiotic selection marker after homologous recombination in *C. jejuni* 11168H to give pMHT. To add homologous recombination arms for *cmeABC*- *aphA*, pMH3 was cut by SaCII to ligate *cj0364* at the 3’ end of *aphA* to give pMHTF. For g0*cmeABC-aphA*, each asparagine in the non-canonical glycosylation sequon (D/E-X_1_-N-X_2_- S/T where X_1_ and X_2_ are any amino acid except proline) was altered to glutamine *in-silico* and nucleotide sequence of g0*cmeABC* was synthesized by (Clonetech, USA) DNA was then treated as above to generate pATKH.

To generate *C. jejuni cmeD*::*cat*, electroporation of pATN into *C. jejuni* 11168H was carried out as previously described^10^. The transformants were selected on CBA plates supplemented with 10 µg/ml chloramphenicol and the double cross over event was confirmed by PCR, this strain was then used as parent strain to generate other mutants. Plasmids pMHT and pATK were electroporated into *C. jejuni cmeD*::*cat* to generate *C. jejuni cmeD*::*cat cmeC*::*cmeC-aphA* and *C. jejuni cmeD*::*cat cmeABC*::*cmeABC*-(N->Q)-*aphA*, respectively. Transformants were selected on CBA plates supplemented with 10 µg/ml chloramphenicol and 30 µg/ml kanamycin and the double cross over event was confirmed by PCR.

### Antibiotic sensitivity test (E-test)

*C. jejuni* 11168H were grown in suspension in Mueller-Hinton broth equivalent to 1.0 MacFarland’s standard and 100 µl aliquots were spread plated on dry Mueller-Hinton agar plates supplemented with 5 % Sheep blood (Oxoid, UK), the plates were left for 5-10 minutes to dry before the antibiotic strip (Oxoid, UK) was added. Plates were incubated at 37°C overnight. The minimum inhibitory concentration (MIC) was read directly from the strip at the oint where the zone of inhibition of bacterial growth intersected with the antibiotic concentration on the strip.

### Ethidium bromide accumulation assay

Bacterial cells were grown to mid log phase (OD _600_ 0.4-0.5). Cells were harvested, washed and resuspended in 0.1M sodium phosphate buffer pH 7 (previously incubated in the VAIN) to OD _600_ 0.2. Cells were then incubated in the VAIN for 15 mins at 37°C o before a 100µl aliquot was withdrawn to indicate time zero. Ethidium bromide (Sigma, UK) was added to final concentration 2 µg/ml and fluorescence was measured at 530 nm excitation and 600 nm emission using a plate reader (Molecular Devices M3 plate reader, USA).

### Expression of CmeA and gCmeA

Protein expression was carried out in *E. coli* strains unless stated otherwise. CmeA and CmeC were expressed in *E. coli* DH10B carrying pMH5 plasmid and pAT3, respectively, whilst gCmeA was expressed in *E. coli* SDB1 carrying pGVXN114, pWA2 and pACYC*(pgl*). Initiating cultures were grown overnight in LB broth supplemented with appropriate antibiotics at 37 °C under shaking condition. The following day, 10 ml of culture was withdrawn from the shake flask to inoculate 400 ml LB broth supplemented with appropriate antibiotics. To achieve optimal glycosylation of CmeA, PglB was expressed from pGVXN114 by the addition of 0.5 mM ITPG at OD _600_ 0.5-0.6. Cultures were incubated at 37°C for 24 hours with shaking. Cultures were centrifuged and cell pellets washed with binding buffer (300 mM NaCl, 50 mM NaH_2_PO_4_ with 25mM imidazole) and passed twice through a high pressure cell homogeniser (Stanstead works, UK). Cell debris was removed by centrifugation at 20,000 xg for 45 minutes. Supernatant was collected and incubated with 0.2 ml Ni-NTA for 1 hour at 4 °C then washed with 50 ml binding buffer and eluted four times in 0.5 ml elution buffer (300 mM NaCl, 50 mM NaH_2_PO_4_ with 250mM imidazole).

### Cloning and expression of CmeC

To express CmeC in *E. coli*, *cmeC* lacking signal peptide sequence was amplified by PCR with CmeCFwd1 and CmeCRev using *C. jejuni* 11168H genomic DNA as a template. The amplicon was then cut by NheI and SalI and cloned into pEC415 downstream of the DsbA signal peptide sequence to give pCMECDSBA. *E. coli* carrying pCMECDSBA was grown in LB media supplemented with ampicillin (100 µg/ml) overnight at 37 °C under shaking condition. On the following day, 10 ml were withdrawn from the overnight culture to inoculate 400 ml LB media. Cells were grown to OD _600_ 0.5-0.6 and 0.2% L-arabinose was added to induce the expression of CmeC. Cultures were incubated at 37 °C for 24 hours with shaking at 180 rpm. Cultures were centrifuged and cell pellets washed with binding buffer (300 mM NaCl, 50 mM NaH_2_PO_4_ with 25mM imidazole) and passed twice through cell homogeniser (Stanstead works, UK). Cells debris was removed by centrifugation at 20,000 xg for 45 minutes and then collected and incubated in binding buffer with 2 % DDM for 3 hours at 4 °C. The mixture was then centrifuged at 15,000 xg for 10 minutes. The supernatant was collected, diluted with binding buffer and incubated with 0.2 ml Ni- NTA for 1 hour at 4 °C then washed with 50 ml binding buffer and eluted four times in 0.5 ml elution buffer (300 mM NaCl, 50 mM NaH_2_PO_4_ with 250mM imidazole).

### CD Spectroscopy

All CD spectra of gCmeA and CmeA were acquired in 0.5mm rectangular cell pathlength using Chirascan spectrometer (Applied Biophysics, UK) equipped with Quantum NorthWest TC125 Peltier unit. Temperature dependent confirmation changes were monitored at wavelength 260-195nm for gCmeA (0.2 mg/ml) and CmeA (0.2 mg/ml) in 10 mM Sodium phosphate, 75 mM Sodium chloride, 10 % glycerol buffer (pH=8.0) during stepwise increase in temperature from 6°C to 94°C. Temperatures were measured directly with a thermocouple probe in the sample solution. Melting temperatures were determined from the derivative CD- Temperature spectra and fitted using a Levenberg–Marquardt algorithm (LMA) on the van*’*t Hoff isochore. (Global 3, Global Analysis for T-ramp Version 1.2 built 1786, Applied Photophysics Ltd, 2007-2012). For Conformation Reversibility Study, far-UV CD spectra were recorded at 20°C, raised to T_m_ and re-cooled to 20°C. The temperature at each elevated T_m_ was kept constant for 5 minutes and the CD spectrum was recorded to assess the rate of protein unfolding process.

### Surface Plasmon Resonance

For coupling of CmeA and gCmeA to the CM5 sensor chip, carboxyl groups on the surface were activated by injecting a 1:1 mixture of 0.4M 1-ethyl-3-(3- dimethylaminopropyl)-carbodiimide (EDC) and 0.1 M N-hydroxysuccinimide (NHS) for 7 minutes at 5 μl/min. CmeA and gCmeA were diluted to 10-20 µg/ml in 0.1 M acetate pH 5.5 and immobilised at 5 μl/min. Immobilisation was stopped when the required RU was achieved. This was followed by injecting 1M ethanolamine pH 8.5 (7 minutes at 5 μl/min) to inactivate excess reactive groups. To account for non-specific binding, a control flow cell was generated using the same method described minus the protein immobilisation step. For coupling of CmeA and gCmeA to a NTA chip, the chip was cleaned and loaded with NiCl_2_ (0.5 mM). The flow cells were then activated as above and CmeA and gCmeA (10 ug/ml in HBSP buffer) were loaded into appropriate flow cells until appropriate RU were achieved. Subsequently the flow cells were treated with ethanolamine as above to block remaining activated sites.

Cmec at various concentrations (3 nM- 0.2 µM) was analysed at a constant temperature of 25°C under continuous flow of HBS-PE buffer (10mM HEPES pH 7.4, 3 mM EDTA, 0.005% (w/v) Surfactant P20 (GE Healthcare) at 30μl/min (sufficient to prevent mass transfer effects) at pH 7.4 for 3 minutes association and a dissociation time of 5 minutes. Experiments at pH 6.0 were performed with 10 mM MES pH 6.0, 3 mM EDTA, 0.005% (w/v) Surfactant P20 (GE Healthcare) The surface chip was regenerated by injecting 0.1 M triethanolamine pH 11.5. Data was analysed using the BIAevaluation software version 4.1.1 (Biacore, GE Healthcare, Amersham). Blank flow cell controls were subtracted. The k_d_ was defined between 10s after the end of the sample injection and 300 sec later.

## Acknowledgements

We acknowledge the Wellcome Trust grant 102978/Z/13/Z for funding.

## Supplementary figures

**Supplementary 1.**
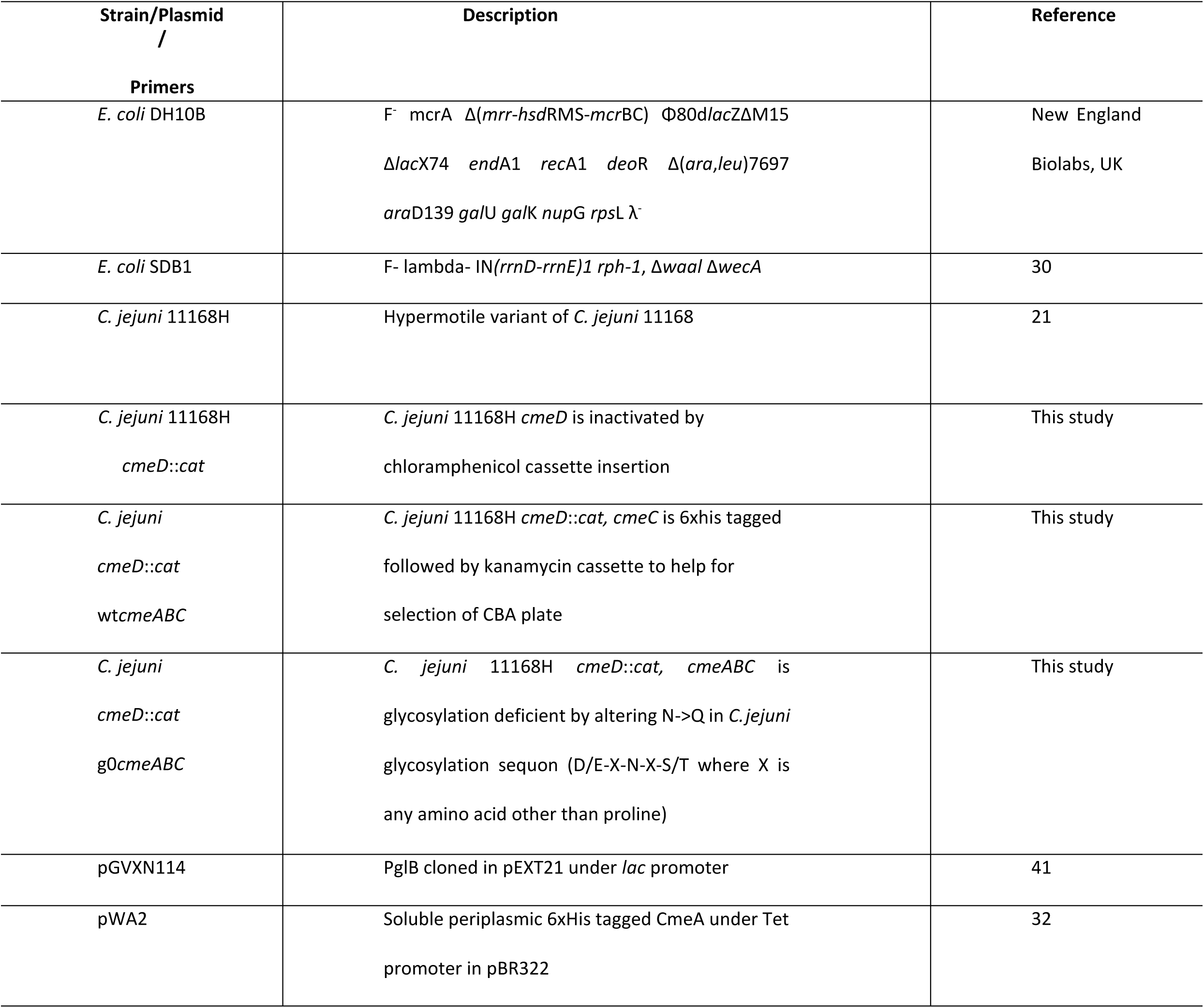
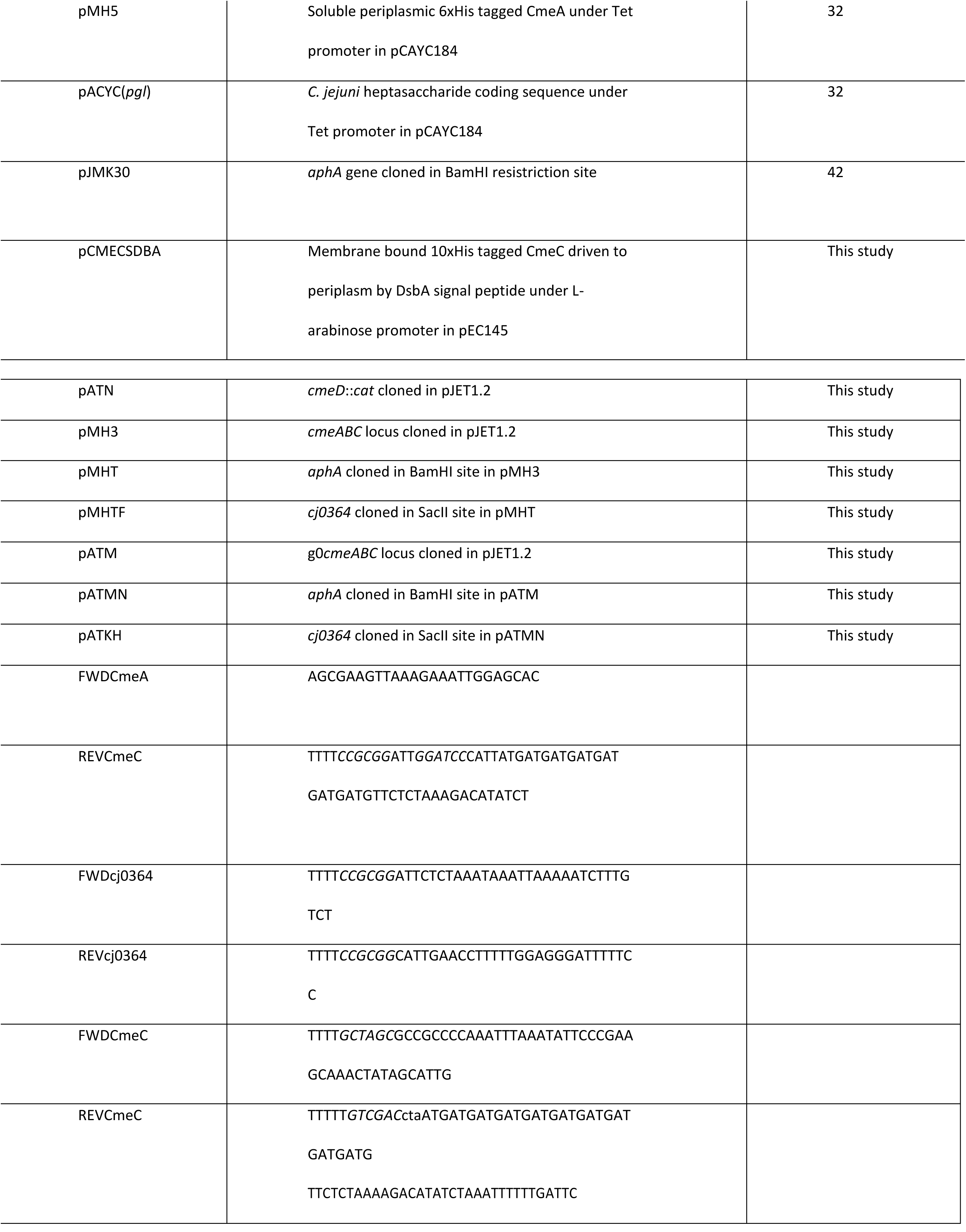
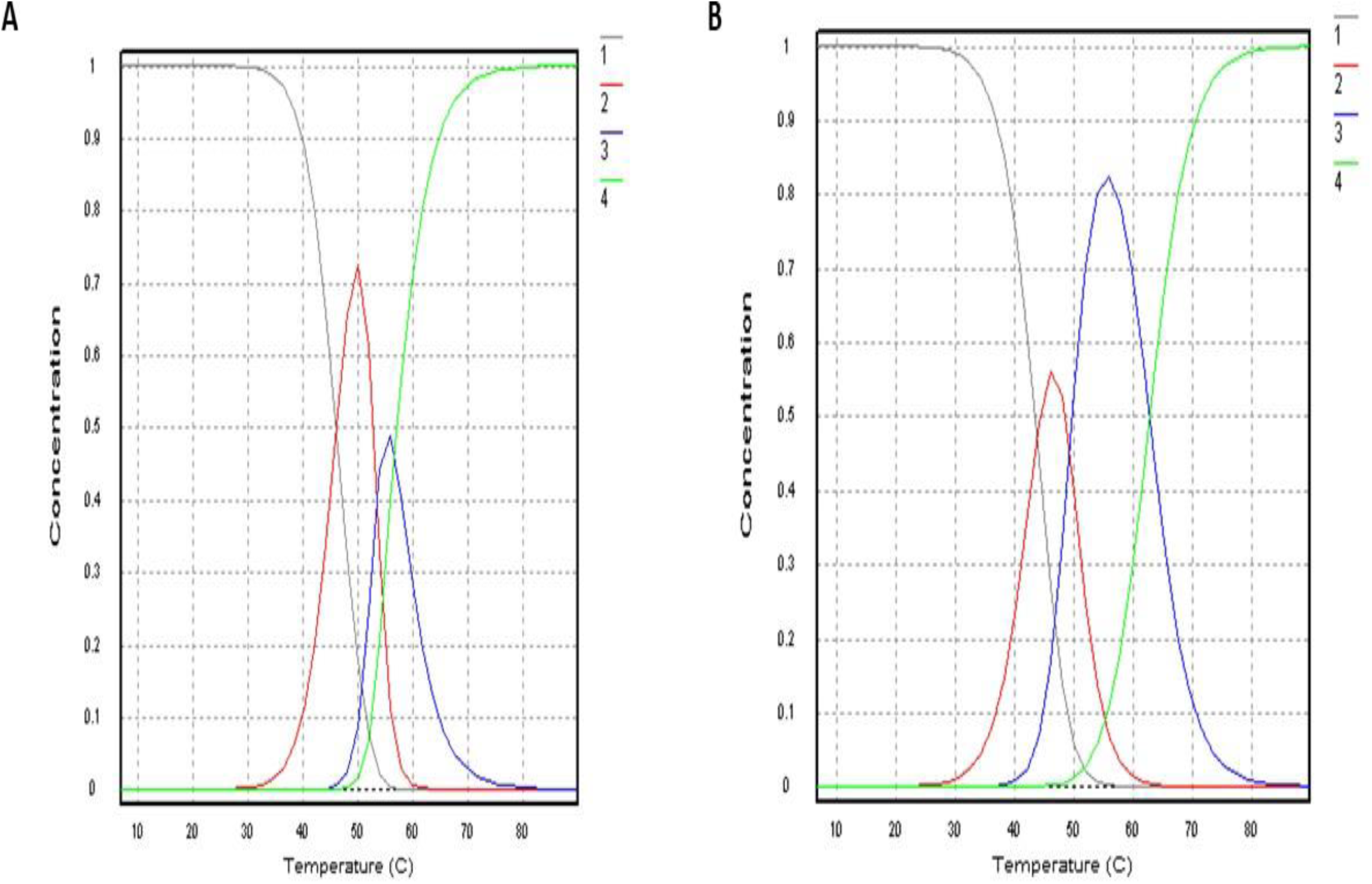
Thermal melts of g0CmeA and g2CmeA in 10 mM sodium phosphate, 75 mM sodium chloride and 10% glycerol (pH 8.0) Concentration as function to temperature representing three transition melting phases for g0CmeA,(A); g2CmeA (B).

**Supplementary 2.**
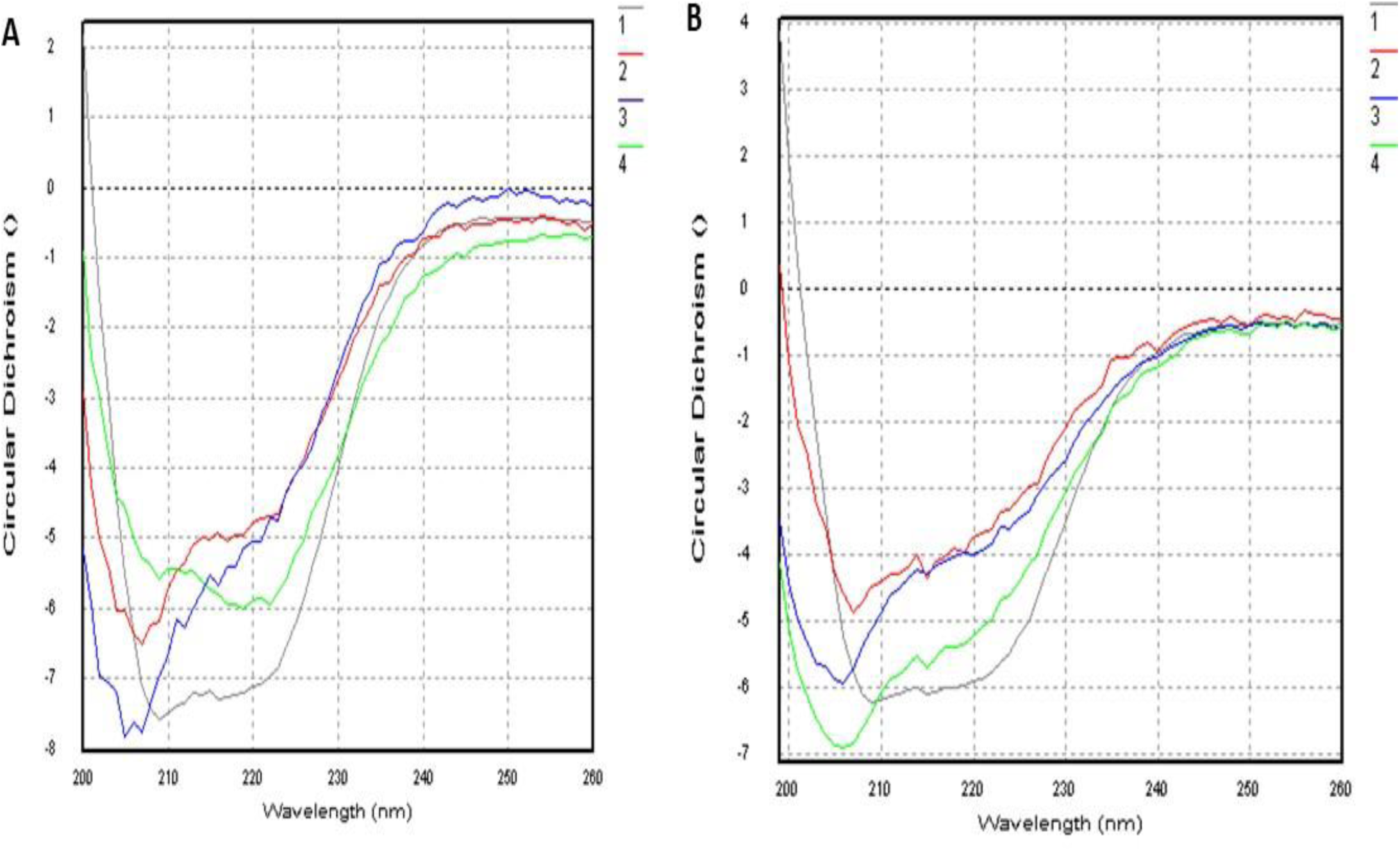
Thermal melts of g0CmeA and g2CmeA in 10 mm sodium phosphate, 75 mM sodium chloride and 10% glycerol (pH 8.0). CD spectra as function to temperature representing more than one melting phases for g0CmeA, (A); g2CmeA, (B).

**Supplementary 3.**
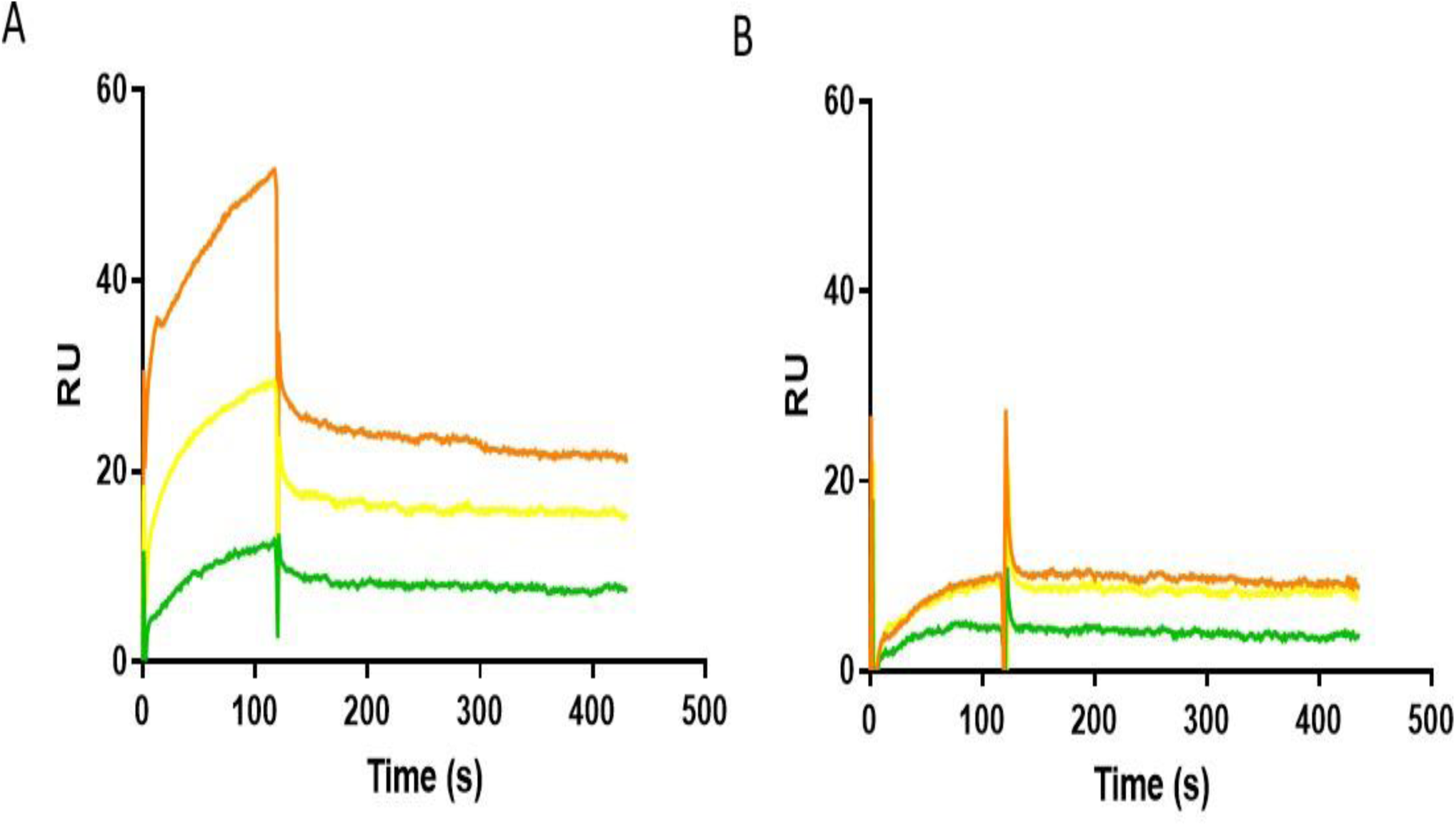
SPR analysis of A) 1090 RU of immobilised g2cmeAand B) 1000RU of immobilisation g0cmeA binding to cmeC offered at 1 ×10^−7^ M (orange); 5 ×10^8^M (yellow) 2.5×10^−8^ M (green) for 2 mins and 5 mins dissociation. CmeA variants were covalently associated by NHS/EDC after association through C-terminal 6Xhis-tag association with the NTA surface.

**Supplementary 4.**
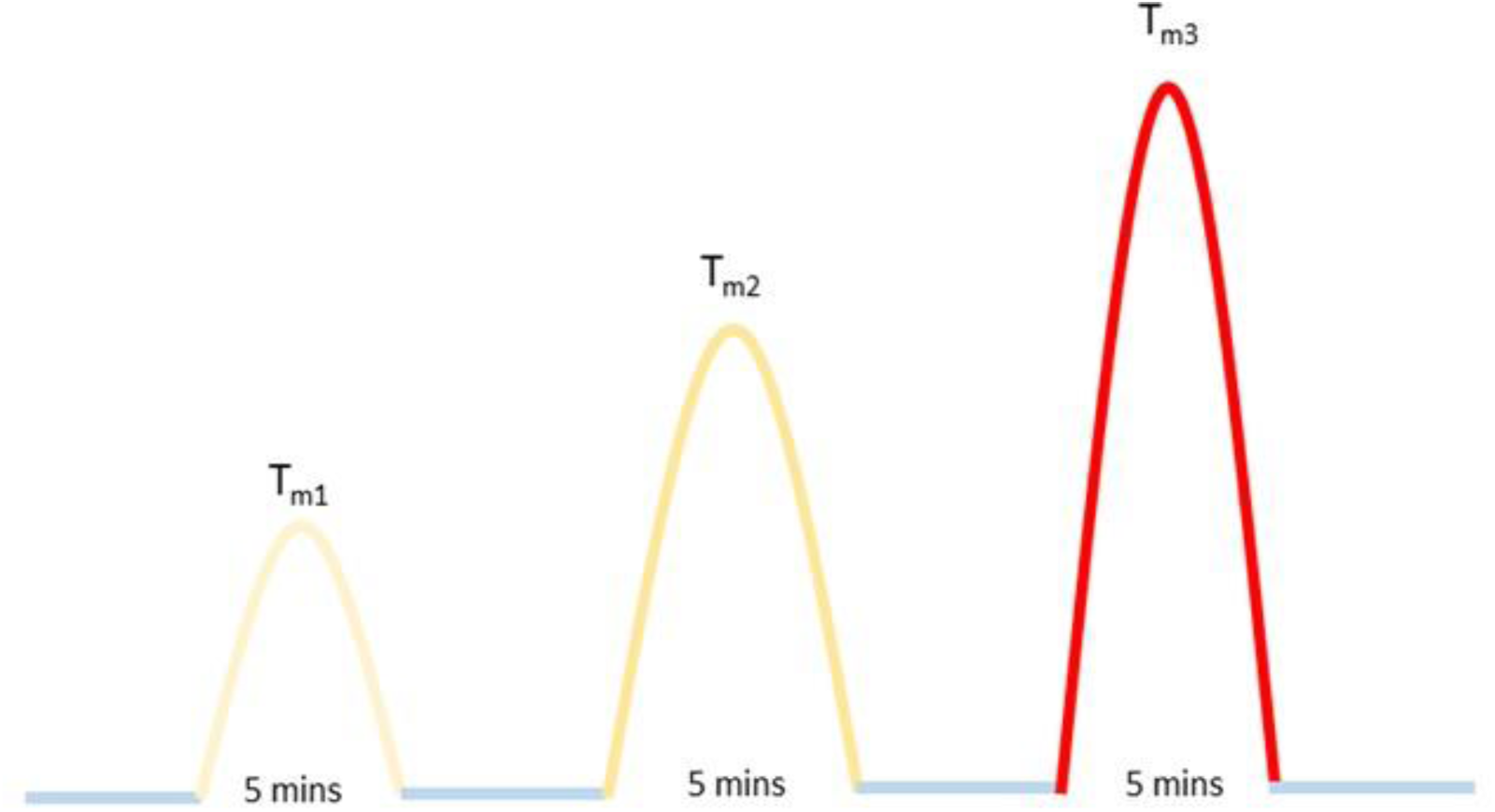
Scheme representing reversibility study CmeA variants were cooled at 20 C (blue) then heated up to T_m_ for 5 minutes then cooled again at 20 C the corresponding T_m_ are shown from golden yellow to red, CD spectra were recorded at each temperature.

